# Dynamic NHE1-Calmodulin complexes of varying stoichiometry and structure regulate Ca^2+^-dependent NHE1 activation

**DOI:** 10.1101/2020.08.04.236463

**Authors:** Lise M. Sjøgaard-Frich, Andreas Prestel, Emilie S. Pedersen, Marc Severin, Johan G. Olsen, Birthe B. Kragelund, Stine F. Pedersen

## Abstract

Calmodulin (CaM) engages in Ca^2+^-dependent interactions with numerous proteins, including human Na^+^/H^+^-exchanger NHE1. Using nuclear magnetic resonance (NMR) spectroscopy, isothermal titration calorimetry, and fibroblasts expressing wildtype and mutant NHE1, we discovered multiple accessible states of this important complex existing in different NHE1:CaM stoichiometries and structures. We solved the NMR solution structure of a ternary complex in which CaM links two NHE1 cytosolic tails. *In vitro*, stoichiometries and affinities were tunable by variations in NHE1:CaM ratio and calcium ([Ca^2+^]) and by phosphorylation of S648 in the first CaM-binding α-helix. In cells, Ca^2+^-CaM-induced NHE1 activity was reduced by mimicking S648 phosphorylation or mutating the first CaM-binding helix, whereas Ca^2+^-induced NHE1 activity was unaffected by inhibition of Akt, one of several kinases phosphorylating S648. Our results reveal the diversity of NHE1:CaM interactions and suggest that CaM may contribute to NHE1 dimerization. We propose that similar structural diversity is relevant to other CaM complexes.

## Introduction

Calmodulin (CaM) is a ubiquitously expressed EF-hand Ca^2+^ -binding hub protein, which regulates a plethora of Ca^2+^ -dependent cellular processes such as ion transport, muscle contraction, proliferation and apoptosis [1–5]. CaM is a small, α-helical protein consisting of two similar but not identical, N- and C-terminal lobes (N-lobe and C-lobe) connected by a helical linker with a disordered, flexible hinge. Upon changes in the free cytosolic Ca^2+^ concentration, [Ca^2+^]_i_, CaM binds up to four Ca^2+^ ions. This induces structural rearrangements exposing hydrophobic patches through which CaM interacts with numerous structurally and functionally different proteins [2,6]. In the vast majority of cases, CaM wraps both its lobes around its α-helix-forming targets, exemplified by the interaction with myosin light chain kinase [7,8]. Although this binding mode has been coined canonical, a single CaM binding motif does not exist. Indeed, in some cases only CaM with Ca^2+^ present in one lobe is recognized, whereas in other cases only the apo-form of CaM is bound [9–11]. Still, there are common features of CaM binding regions, such as high α-helix propensity, net positive charge, and obligate hydrophobic docking residues [2]. Examples include the CaM substrates estrogen receptor alfa (ERα) [12,13], the voltage-gated K^+^ channel Kv10.1 (eag1) [14,15], and aquaporin 0 [16].

One widely studied CaM-binding protein is the Na^+^/H^+^ exchanger, NHE1 (SLC9A1), a major acid-extruding transporter in essentially all mammalian cells studied [17,18]. Upon activation by cytosolic acidification or a wide range of mitogenic and other signaling events, NHE1 extrudes H^+^ from the cytosol in exchange for Na+ ions. In addition to its role in intracellular pH (pH_i_) regulation, NHE1 regulates key cellular functions such as proliferation, growth/death balance, cell volume, cell motility, and tissue homeostasis [17,18]. Consequently, NHE1 hyperactivity is implicated in many important pathologies including cancers [19,20], as well as cardiovascular disease, especially ischemia-reperfusion damage and cardiac hypertrophy [21–23] and non-alcoholic steatohepatitis (NASH) [24].

NHE1 consists of a 12-transmembrane (TM)-helix transport domain and a ~300 residue long cytoplasmic C-terminal domain, which has extensive disordered regions [25]. *Via* this tail, NHE1 activity is regulated through multiple phosphorylation-dephosphorylation events and interactions with a plethora of binding partners including CaM (for reviews, see [17,18,26]). CaM binds directly to the cytoplasmic domain of NHE1 in the presence of Ca^2+^ [27,28] as originally demonstrated *in vitro* by binding of either an NHE1 fusion protein or full-length protein [28]. Two neighboring α-helical CaM binding sites of NHE1 each bind CaM, but with different affinities. For an NHE1-variant deleted in CaM-binding region 2 (CB2, defined as D656-L691), *K*_*d*_ was ~20 nM and for a NHE1-variant deleted in CaM-binding region 1 (CB1, defined as N637-A656) *K*_*d*_ was ~350 nM (Fig. 1a-b) [28]. A crystal structure of CaM bound to a peptide of NHE1 containing both CaM binding sites (A622-R690) revealed an unusual binding mode. CaM assumed an extended configuration, and the N-terminal CaM binding site of NHE1 (H1 in the following) made an unusual interaction with the "back side" of the C-lobe, whereas the N-lobe of CaM engaged with the low affinity C-terminal site of NHE1 (H2 in the following) forming an α-helix in the hydrophobic ligand-binding groove [27].

**Fig 1.**
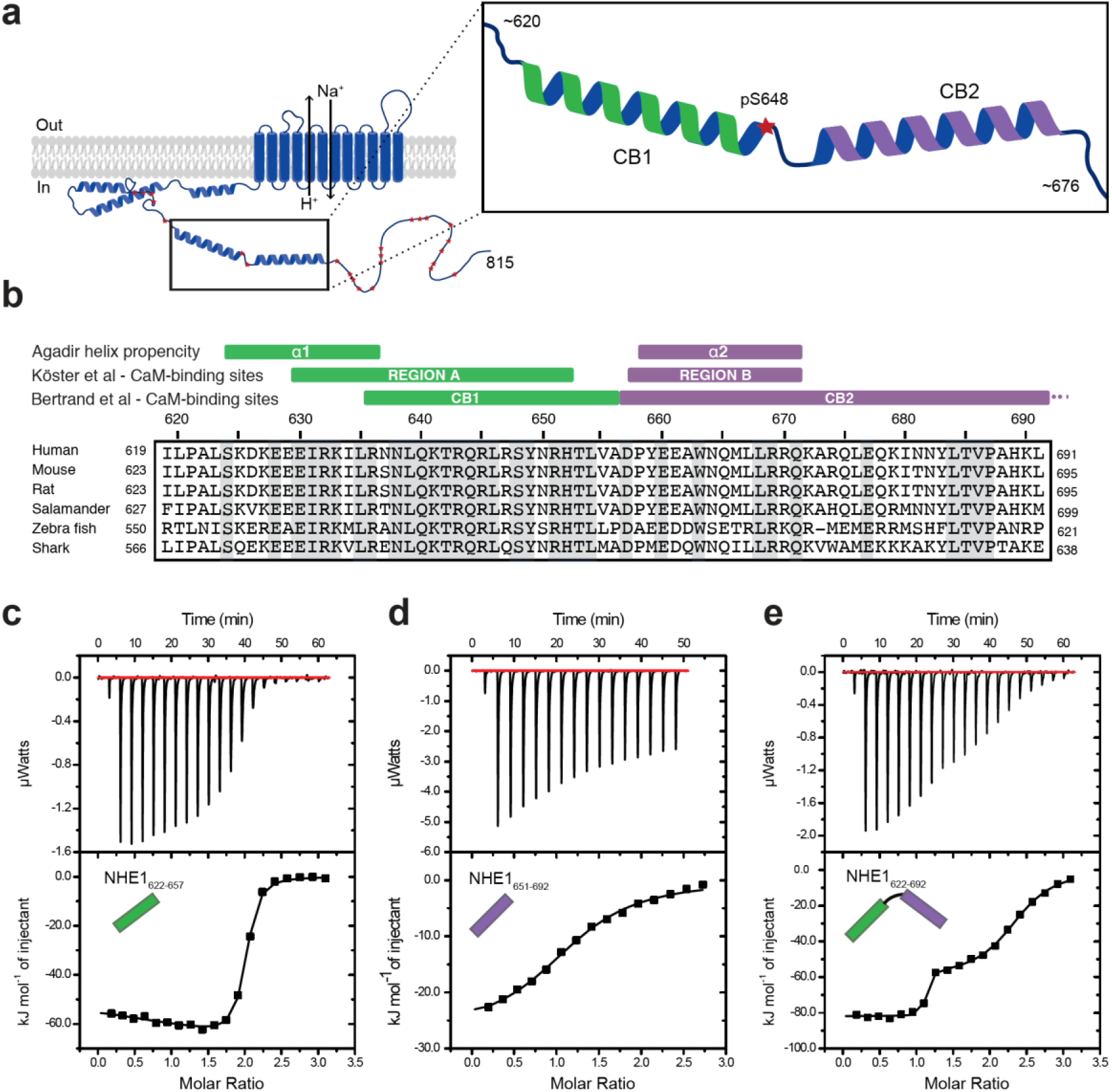
*NHE1:CaM interactions* in vitro *adopt different stoichiometries*. (a) Sketch of NHE1 (blue) in the membrane highlighting helical elements in the cytoplasmic tail and known phosphorylation sites (red stars). Magnified region depicts the two previously described CaM-binding (CB) α-helices CB1 (green) and CB2 (purple) with the phosphorylated S648. (b) Alignment of the sequences of human, mouse, rat, salamander, zebra fish and shark NHE1 highlighting in grey identical residues. Indicated above are structural and functional regions based on either an Agadir α-helix prediction, the CaM binding Region A (green) and Region B (purple), as defined by Koster et al. [27] or the CaM-binding region I (CB1) and II (CB2), as defined by Bertrand et al [28]. Binding NHE1 to CaM by ITC. In representative ITC experiments we injected (c) NHE1_622-657_ or (d) NHE1_651-692_ or (e) NHE1_622-692_ into CaM. The upper parts show baseline-corrected raw data from the titrations and the lower parts the normalized integrated binding isotherms with the fitted binding curves assuming a single binding event in (d), and two binding events in (c) and (e). The NHE1 peptides titrated into CaM are shown in cartoon. The ITC-raw data and the experiments not shown are available in **Source data file** 1.1-1.13.

Binding of Ca^2+^-loaded CaM to NHE1 in response to an increase in [Ca^2+^]_i_ elicits an alkaline shift in the pH_i_ sensitivity of NHE1 in cells, resulting in its activation at less acidic pH_i_. Furthermore, deletion or mutation of CB1 results in loss of such [Ca^2+^]_i_-responsiveness [28,29]. Mechanistically, NHE1:CaM interaction has been proposed to activate NHE1 by relieving an autoinhibitory interaction of the C-terminal tail of NHE1 with its TM region [30,31] and has been implicated in activation of NHE1 by a wide range of hormones and growth factors, including epidermal growth factor, serotonin, angiotensin and endothelin [32–34]. NHE1:CaM interaction was suggested to be regulated by Akt kinase-mediated phosphorylation of S648 of human NHE1 [35], which is localized at the C-terminal distal part of the first CaM-binding helix. However, the role of S648 phosphorylation in NHE1 regulation has been reported both as inhibitory [35] and stimulatory [36], pointing to a possible context-dependence of NHE1:CaM complex formation.

These observations led us to speculate that the NHE1:CaM complex exhibits a greater degree of diversity than revealed by crystallographic analyses. To explore this, and to elucidate the precise mechanism and role of S648 phosphorylation, we employed a combination of solution state-nuclear magnetic resonance (NMR) spectroscopy, isothermal titration calorimetry (ITC), NHE1 mutations, proximity ligation assays (PLA), and real-time analyses of NHE1-mediated acid extrusion after stable expression of wild type and variant NHE1 in mammalian fibroblasts. Our findings reveal that in solution, CaM interacts with NHE1 by exploiting different binding modes and stoichiometries, thereby forming several possible structures. Accordingly, in cells, Ca^2+^-induced NHE1 activation, but not NHE1:CaM proximity, was modulated by mutations preventing (S648A) or mimicking (S648D) phosphorylation, as well as reduced by charge-reversal mutations of the first CaM-binding site. This was not dependent on Akt kinase, inhibition of which did not affect Ca^2+^-induced activation of NHE1 under these conditions. Consistent with this notion, S648 phosphorylation could be mediated by other Ser/Thr kinases. We propose that the interaction between CaM and NHE1 is highly dynamic, involving both CaM binding helices of NHE1, and with a functional role of S648 that is not exclusively downstream from Akt. We suggest that the high degree of diversity observed for NHE1:CaM complexes may be relevant to other CaM complexes.

## Results

### CaM interacts with NHE1 in multiple conformations *in vitro*

In order to characterize the interaction of CaM with NHE1 *in vitro*, we designed three NHE1 peptides: human (h)NHE1_622-657_ (H1), hNHE1_651-692_ (H2) and hNHE1_622-692_ (H1H2); all based on previous studies [27], sequence conservation and helix propensity calculations (Fig. 1a-b). First, ITC was applied to determine binding thermodynamics and stoichiometries of the NHE1-derived peptides and CaM. In line with previous results [28], H1 bound CaM with high affinity. In addition, we observed that one CaM molecule bound two H1 peptides (n = 1.92 ± 0.06, Fig. 1c); an interaction mode not expected from the crystal structure [27]. Fitting the ITC data to a binding model with two independent sites revealed similar high affinities (*K*_*d,1*_ = 27 ± 8 nM; *K*_*d,2*_ = 42 ± 6 nM, Fig. 1c), but with different enthalpic and entropic contributions (Table 1). The affinity of the second helix, H2, for CaM was almost three magnitudes weaker (*K*_*d*_ = 9.0 ± 0.6 μM) than that of H1. This interaction showed only one transition, indicating the formation of a 1:1 complex (n = 1.15 ± 0.08, Fig. 1d). Finally, the binding isotherm for the H1H2 peptide displayed two distinct transitions, indicating successive binding events and/or diverse binding modes with different CaM:H1H2 stoichiometries. Fitting to a two-site binding model allowed us to determine the affinity of the first binding event to *K*_*d,1*_ = 0.27 ± 0.07 nM with a stoichiometry of one (n_1_ = 1.04 ± 0.05), and the affinity of the second binding event to *K*_*d,2*_ = 560 ± 15 nM with n_2_ = 1.22 ± 0.06, summing up to a total stoichiometry of 1:2 (Fig. 1e).

**Table 1.** Thermodynamic parameters of NHE1:CaM interactions.

| | $\Delta H_1$<br>(kJ/mol) | $T\Delta S_1$<br>(kJ/mol) | $K_{d,1}$<br>(nM) | $n_1$ | $\Delta H_2$<br>(kJ/mol) | $T\Delta S_2$<br>(kJ/mol) | $K_{d,2}$<br>(nM) | $n_2$ |
| --- | --- | --- | --- | --- | --- | --- | --- | --- |
| **H1**<br>**(one site)** | $-60 \pm 3$ | $-17 \pm 2$ | $23 \pm 5$ | $1.92 \pm 0.06$ | -- | -- | -- | -- |
| **H1**<br>**(two site)** | $-39 \pm 11$ | $4 \pm 10$ | $27 \pm 8$ | $0.92 \pm 0.05$ | $-83 \pm 8$ | $-41 \pm 8$ | $42 \pm 6$ | $0.98 \pm 0.01$ |
| **H2**<br>**(one site)** | $-29 \pm 4$ | $0 \pm 3$ | $8900 \pm 600$ | $1.15 \pm 0.08$ | -- | -- | -- | -- |
| **H1H2**<br>**(two site)** | $-84 \pm 5$ | $-29 \pm 5$ | $0.27 \pm 0.07$ | $1.04 \pm 0.05$ | $-62 \pm 3$ | $-26 \pm 3$ | $560 \pm 10$ | $1.22 \pm 0.06$ |
| **H1H2 pS648**<br>**(two site)** | $-75 \pm 3$ | $-27 \pm 2$ | $5 \pm 4$ | $0.93 \pm 0.02$ | $-58 \pm 7$ | $-26 \pm 8$ | $2700 \pm 700$ | $1.16 \pm 0.06$ |

**Table 2.** Exchange kinetics from 2D NMR lineshape analysis.

| | $k_{on,1}$ ($M^{-1}s^{-1}$) | $k_{off,1}$ ($s^{-1}$) | $K_{d,1}$ (nM) | $k_{on,2}$ ($M^{-1}s^{-1}$) | $k_{off,2}$ ($s^{-1}$) | $K_{d,2}$ (nM) |
| --- | --- | --- | --- | --- | --- | --- |
| **H1**<br>**(two site)** | $(1.5 \pm 0.6) \times 10^9$ | $40 \pm 5$ | $(27)^{\#}$ | $(3.7 \pm 1.3) \times 10^8$ | $16 \pm 3$ | $(42)^{\#}$ |
| **H2**<br>**(one site)** | $(1.6 \pm 0.1) \times 10^8$ | $2600 \pm 100$ | $16000 \pm 1000$ | -- | -- | -- |

<sup>#</sup> $K_{ds}$ taken from ITC analysis, kept constant during fitting

These results demonstrate formation of variable NHE1:CaM complexes *in vitro* and show that the interaction cannot be explained by a simple 1:1 complex. Various complexes with different stoichiometries as well as thermodynamic profiles are formed depending on the binding site availability and NHE1:CaM ratio.

### Both lobes of CaM interact independently with H1 with high affinity

To gain structural insight into these phenomena, we used solution NMR spectroscopy. We titrated ^15^N-labelled CaM with increasing concentrations of unlabelled H1 and followed the changes in shape, intensity and position of the NMR peaks (Fig. 2a). Upon addition of H1 (50 μM CaM + 50 μM NHE1), most signals from both CaM lobes broadened drastically, indicating exchange between free and bound state(s) on an intermediate NMR time scale (*k*_*ex*_ *≈ Δδ*) [37] (Fig. 2b). This suggests dynamics in the complex between CaM and H1 occurring on the μs to ms timescale. After addition of excess H1 peptide (50 μM CaM + 125 μM H1, denoted 1:2 in the following), the signals re-sharpened and no spectral changes were observed upon further addition of H1 (Fig. 2a-b). This is in line with the CaM:H1 stoichiometry of 1:2 obtained by ITC. At a molar ratio of 1:1, all signals were broadened by exchange processes, but it was evident that the main signal intensities from the C-terminal lobe of CaM were located close to the position of the fully bound state, while signals of the N-lobe were positioned close to those of the free form of CaM, as exemplified for G135 and G62, respectively (Fig. 2b). This indicated that in the main state at a 1:1 stoichiometry, H1 is bound to the higher affinity C-lobe, as depicted in the cartoon in Fig. 2c. Not all peaks could be followed throughout the titration, but the high-quality NMR spectra of the 1:2 (CaM:H1) complex allowed assignments of CaM in the ternary complex and mapping of the chemical shift perturbations (CSPs) onto the sequence (Fig. 2d). Large CSPs were observed in both lobes and for residues throughout CaM, highlighting binding of H1 to both lobes. The dissociation rate of H1 from the C-lobe and N-lobe was determined by 2D NMR lineshape analysis to *k*_*off,1*_ = 40 ± 5 s^−1^ and *k*_*off,2*_ = 16 ± 3 s^−1^, respectively (Figure 2 – Figure supplement 1a), revealing a >10 times faster association with the C-lobe than with the N-lobe. To further substantiate these results, the opposite titration was performed: ^15^N-labelled H1 was monitored as the concentration of unlabelled CaM was increased. Only a few signals from the free peptide were visible, while at a 1:2 (CaM:H1) molar ratio, two sets of signals became distinct and could be assigned to H1 bound to either of the two CaM lobes, respectively (Figure supplement 2a). When more CaM was added, the set of signals originating from H1 bound to the N-lobe decreased in intensity and vanished at a molar ratio of 3:1 (CaM:H1). In this condition, the peptide was almost exclusively bound to the higher affinity C-lobe of CaM (Figure 2 – Figure supplement 2a).

**Fig 2.**
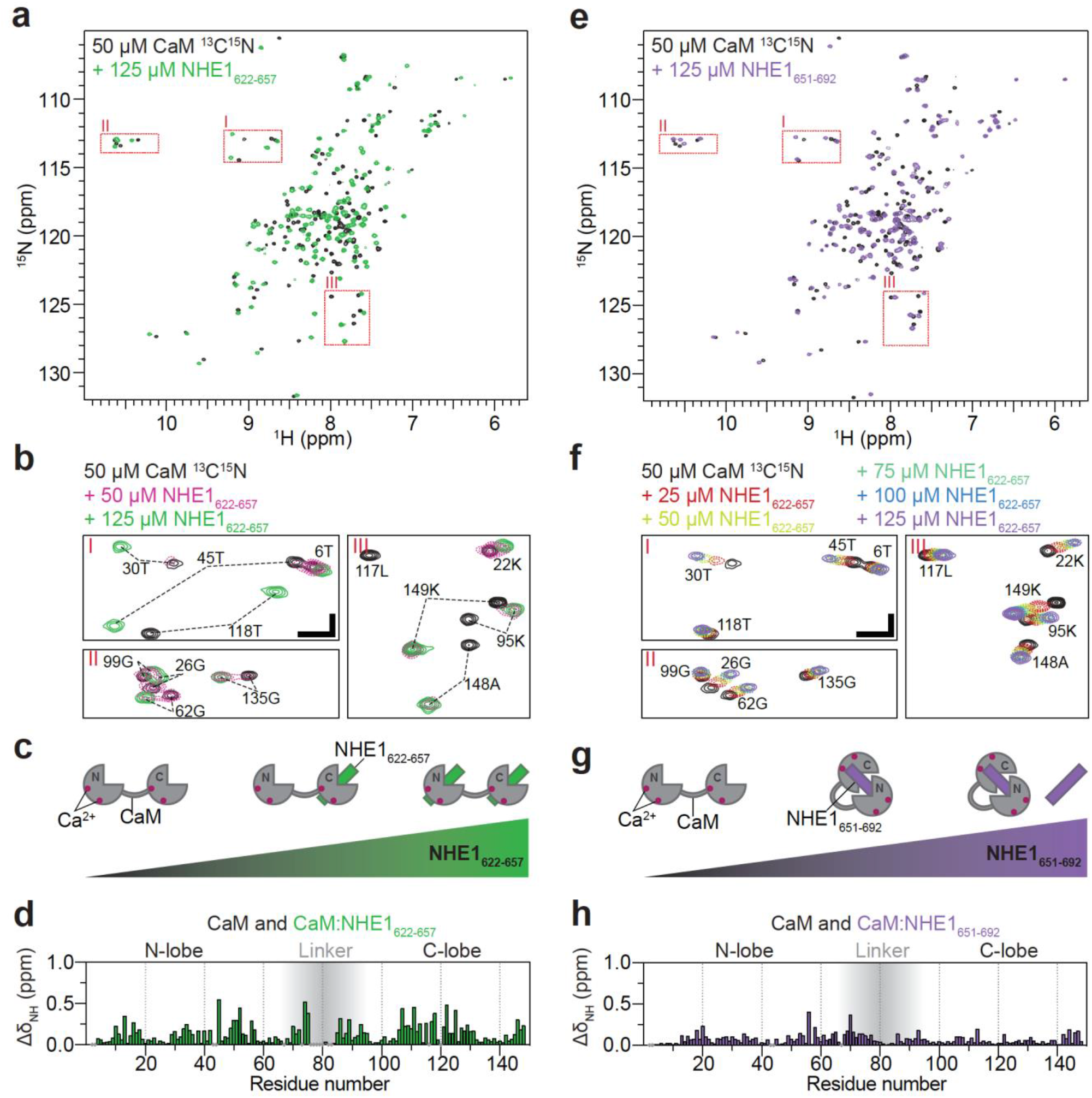
CaM interaction with NHE1_622-657_ (H1) and NHE1_651-692_ (H2) (a) ^1^H, ^15^N HSQC spectra of 50 μM ^15^N-CaM alone (black) and in the presence of 125 μM unlabelled NHE1_622-657_ (H1, green). (b) Zoom on the three highlighted regions from the spectrum in (a); ^1^H,^15^N HSQC spectra of 50 μM ^15^N-CaM in the presence of 0 μM (black) 50 μM (dashed red) and 125 μM (green) unlabelled NHE1_622-657_. The horizontal and vertical scalebars correspond to 0.1 ppm δ^1^H and 0.5 ppm δ^15^N respectively. (c) Cartoon representation of CaM interaction with increasing amounts of NHE1_622-657_. (d) Chemical shift perturbations (CSP; ΔδNH) of CaM upon interaction with NHE1_622-657_ at saturation (125 μM NHE1_622-657_, 50 μM ^15^N-CaM, denoted 2:1 NHE1:CaM). (e) ^1^H,^15^N HSQC spectra of 50 μM ^15^N-CaM alone (black) and in the presence (purple) of unlabelled NHE1_651-692_. (f) Zoom on the three highlighted regions from the spectrum in (e); ^1^H,^15^N HSQC spectra of ^15^N-CaM titrated from zero to 125 μM (black to purple) with unlabelled NHE1_651-692_ (H2). The horizontal and vertical scalebars correspond to 0.1 ppm δ^1^H and 0.5 ppm δ^15^N respectively. (g) Cartoon representation of CaM interaction with increasing amounts of NHE1_651-692_. (h) CSP (ΔδNH) of CaM upon interaction with NHE1_651-692_ at saturation (125 μM NHE1_651-692_, 50 μM ^15^N-CaM). **Figure supplement 1**: TITAN 2D-NMR lineshape analysis for extraction of binding kinetics. **Figure supplement 2**: Titration of ^15^N labeled H1, H2 and H1H2 with unlabeled CaM. **Figure supplement 3:** Concentration dependence of the NMR chemical shift of NHE1 H2

Titration of ^15^N-labelled CaM with the low affinity H2 peptide resulted in signals gradually moving with increasing peptide concentration, indicating the free and bound states to be in fast exchange on the NMR timescale (*k*_*ex*_ > *Δδ*, Fig. 2e-f) [37]. The change in peak position could be followed throughout the titration, and 2D NMR lineshape analysis resulted in affinity and dissociation rates of *K*_*d*_ = 16 ± 1 μM and *k*_*off*_ = 2600 ± 100 s^−1^, respectively (Figure 2 – Figure supplement 1b). Again, the CSPs spanned both CaM lobes (Fig. 2h), although they were less pronounced for the C-lobe and generally smaller compared to those induced by H1 binding. Inverting the titration gave a similar result (Figure 2 - Figure supplement 2b). However, in this case the spectral quality of ^15^N-labelled H2 in both free and bound states was inferior with very heterogeneous signal intensities and only few assignable signals, likely originating from weak self-association (Figure 2 - Figure supplement 3). This will interfere with CaM interaction and provides a likely explanation for the affinity variations obtained from different titration setups.

Taken together, these data show that each of the CaM lobes can bind one NHE1 H1 with high affinity and that the C-lobe is energetically slightly favored. NHE1 H2 binds with lower affinity in a 1:1 complex with both CaM lobes involved, indicating canonical CaM binding as depicted in the cartoon in Fig. 2g.

### CaM can interact with NHE1 H1H2 in different conformations with different stoichiometries

The two distinct transitions in the ITC isotherm of CaM titration with H1H2 (Fig. 1e) indicated a more complex interaction between CaM and the full CaM binding region of NHE1 compared to each binding site alone. To decipher this using NMR, we titrated unlabelled H1H2 into ^15^N-labelled CaM, and compared the ^15^N-HSQC fingerprint spectra at different stoichiometries to those obtained with H1 or H2 alone. At a molar ratio of 1:2 (CaM:H1H2), the ^15^N-HSQC spectrum closely resembled the state where only H1 is present in 2-fold excess (Fig. 3a-c). This indicated that at this stoichiometry, each CaM lobe interacts with one H1 of H1H2 in a ternary complex, while H2 is not involved in the interaction, as depicted in Fig. 3a. At a stoichiometry of 1:1, the signals of the N-lobe of CaM still overlaid almost perfectly with the spectrum of ^15^N-labelled CaM at saturation with H1. However, signals of the C-lobe now overlaid with the spectrum of CaM bound to H2 (Fig. 3d-f). Thus, at this molar ratio, the major state is the N-lobe of CaM bound to H1 and the C-lobe bound to H2, as depicted in Fig. 3d. A titration of ^15^N-labelled H1H2 with CaM supports these findings, although the spectral analysis was complicated by severe signal overlap (Figure 2 – Figure supplement 2c-d). At a stoichiometry of 1:2 (CaM:H1H2), two sets of signals originated from each residue of H1, while signals from H2 closely resembled those of free H2 (Figure 2 – Figure supplement 2c). This means that H1 was bound to both CaM lobes, while H2 did not partake in the interaction. At a stoichiometry of 1:1, signals from H1 overlapped with the N-lobe bound state, while signals from H2 were invisible or closely resembled the C-lobe bound state (Figure 2 – Figure supplement 2d). For both H1H2:CaM stoichiometries analyzed, the described complexes were the major states as judged by the relative NMR signal intensities. It is, however, important to note that at 1:1 as well as 1:2 (CaM:H1H2), the NMR signals were drastically broadened compared to those of free CaM or of the 1:2 complex saturated with H1 (Fig. 3a,d), indicating a dynamic equilibrium of states with different stoichiometries. Higher order complexes are also possible.

**Fig 3.**
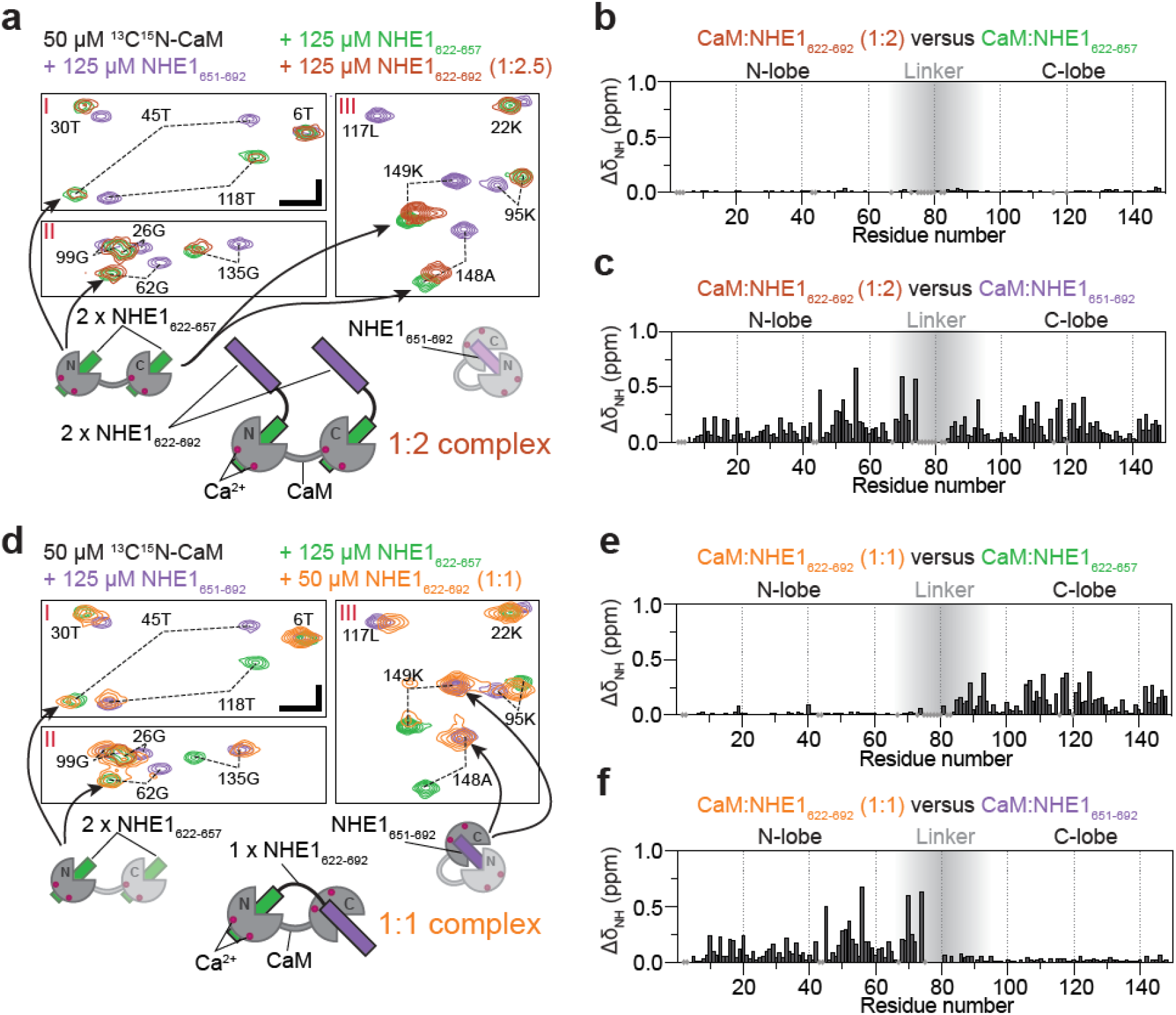
CaM interaction with NHE1_622-692_ (H1H2) at two different stoichiometries. (a) The 1:2 CaM:NHE1_622-692_ (H1H2) complex (molar ratio 1:2). Three regions of the ^1^H,^15^N HSQC spectra of 50 μM ^15^N-CaM with 125 μM NHE1_622-657_ (green), 125 μM NHE1_651-692_ (purple) or 125 μM NHE1_622-692_ (orange) are shown. The horizontal and vertical scalebars correspond to 0.1 ppm δ^1^H and 0.5 ppm δ^15^N respectively. The complexes giving rise to the indicated peaks are depicted in the cartoons below (right and left), with the 1:2 complex in the center. (b) CSPs (ΔδNH) between the 1:2 CaM:NHE1_622-692_ complex and the 1:2 CaM:NHE1_622-657_ complex (dark red and green spectra in (a)). The grey shade illustrates the linker-region of CaM. (c) CSPs (ΔδNH) between 1:2 CaM:NHE1_622-692_ and 1:1 CaM:NHE1_651-692_ (dark red and purple spectra in (a). (d) The 1:1 CaM:NHE1_622-692_ complex. Three regions of the 1H,15N HSQC spectra of 50 μM ^15^N-CaM in the presence of 125 μM NHE1_622-657_ (green), 125 μM NHE1_6521-692_ (purple) or 50 μM NHE1_622-692_ (orange). The horizontal and vertical scalebars correspond to 0.1 ppm δ^1^H and 0.5 ppm δ^15^N respectively. Smaller cartoons represent the 1:2 CaM:NHE1_622-657_ complex (left) and 1:1 CaM:NHE1_651-692_ complex (right) and the 1:1 CaM:NHE1_622-692_ complex identified from overlapping peaks in the ^1^H,^15^N HSQC spectra. (e) CSPs (ΔδNH) between 1:1 CaM:NHE1_622-692_ and 1:2 CaM:NHE1_622-657_ (orange and green spectrum in (d)). (f) CSPs (ΔδNH) between 1:1 CaM:NHE1_622-692_) and 1:1 CaM:NHE1_651-692_ (orange and purple spectrum in (d). The chemical shifts are available in BMRB accession code 34521.

These results show that *in vitro*, NHE1 and CaM can form multiple high affinity complexes of different stoichiometries and structures, depending on the availability of each binding partner.

### Influence of Ca^2+^ availability on complex formation

Cellular [Ca^2+^]_i_ is tightly regulated and numerous Ca^2+^-binding proteins compete for the available pool. The experiments described so far were all performed under Ca^2+^-saturation. To investigate the effect of limited Ca^2+^ availability on the structural ensemble of NHE1 and CaM, we monitored the state of CaM based on the ^1^H-^15^N-HSQC fingerprint spectrum both under nominally Ca^2+^-free conditions and in the presence of the Ca^2+^ chelator ethylenediaminetetraacetic acid (EDTA). In the absence of Ca^2+^, we still observed CSPs of CaM after addition of 2.5-fold molar excess of H1H2 (Supplementary file 1a) suggesting that also the Ca^2+^-free form of CaM binds NHE1 H1H2. Under conditions where both Ca^2+^ (here 2 mM) and high EDTA (here 6.6 mM) are present, free CaM gets stripped of Ca^2+^. However, in the additional presence of 2.5-fold molar excess of H1H2, the signals of the C-lobe overlapped with those from Ca^2+^-CaM bound to H1, while signals of the N-lobe closely resembled the Ca^2+^-free state (Supplementary file 1b). Thus, the presence of H1 increased the affinity for Ca^2+^ in the C-lobe. This further suggests that competition for Ca^2+^ favors another mode of interaction where the CaM C-lobe is bound to Ca^2+^ and to H1 of NHE1, while the N-lobe is in its Ca^2+^-free form.

### Structure of the ternary complex of CaM and two NHE1 H1 helices

Our results suggest that CaM can act as an NHE1 dimerization switch at high [Ca^2+^]_i_, where each of the CaM lobes bind H1, bridging two NHE1 tails. This would provide a structural understanding of previous reports of dimerization sites in the C-terminal tail of NHE1 [38,39], as well as reports based on kinetic analyses of transport indicating that dimerization contributes to the regulation of NHE1 activity [40–43]. To obtain a more detailed understanding of this interaction, we solved the structure of the ternary complex of CaM bound to two H1 peptides using NMR. The results of the structure determination are shown in Fig. 4 and Table 3. Both CaM lobes have one H1 bound inside the hydrophobic cleft (Fig. 4a). An overlay of the 10 lowest energy structures shows that the individual lobes are well defined (Backbone RMSD: N-lobe+H1_N_ = 0.77 ± 0.16 Å; Backbone RMSD C-lobe+H1_c_ = 0.83 ± 0.18 Å), while no NOEs between the two lobes could be obtained, indicating that the linker region remains flexible in the bound state (Fig. 4b-c). In both lobes, the interaction surface is defined by contacts between the hydrophobic face of the amphipathic H1 (L623, I631, I634, L635, N638, L639, T642, L646) (Fig. 4d-e) and the central methionines in the hydrophobic cleft of each CaM-lobe. For the N-lobe this involves M37, M52, M72, M73 and for the C-lobe M110, M125, M145, M146. In addition, a high complementarity in electrostatics is evident, with multiple positively charged residues on the opposite side of NHE1 H1 (K625, K637, R632, K633, R636, K641, R643, R645) complementing the negatively charged surface of CaM (Fig. 4f-i). Although this complex is very different from that shown in the published crystal structure [27], inspection of the neighboring molecules in the crystal lattice reveals that the mode of interaction between the CaM C-lobe and H1, which we describe here by NMR, was also present in the crystal, but was translated as crystal contacts between symmetry-related molecules (Figure 4 – Figure supplement 1).

**Table 3.**
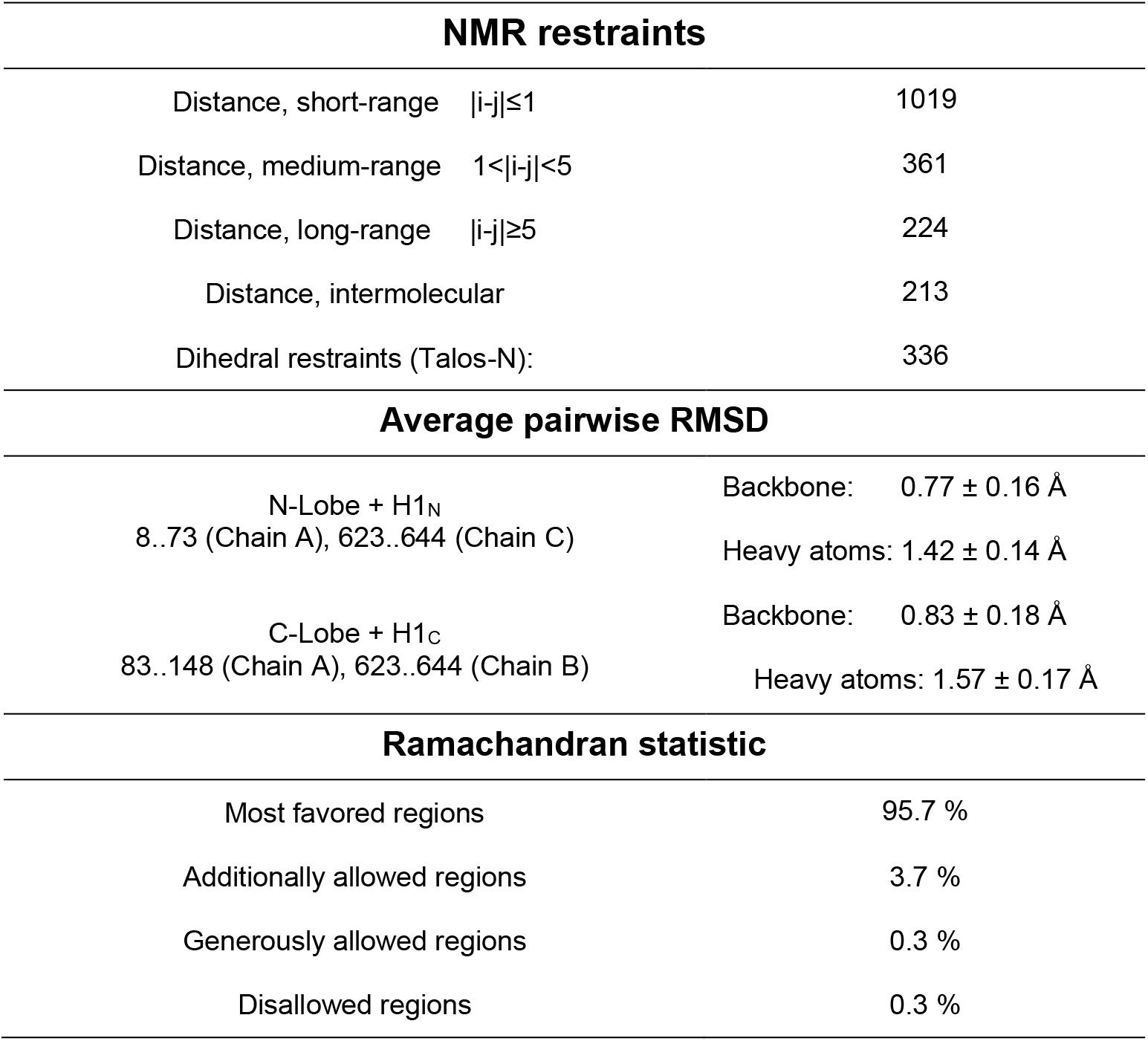
Structural statistics (pdb accession code 6zbi)

**Fig 4.**
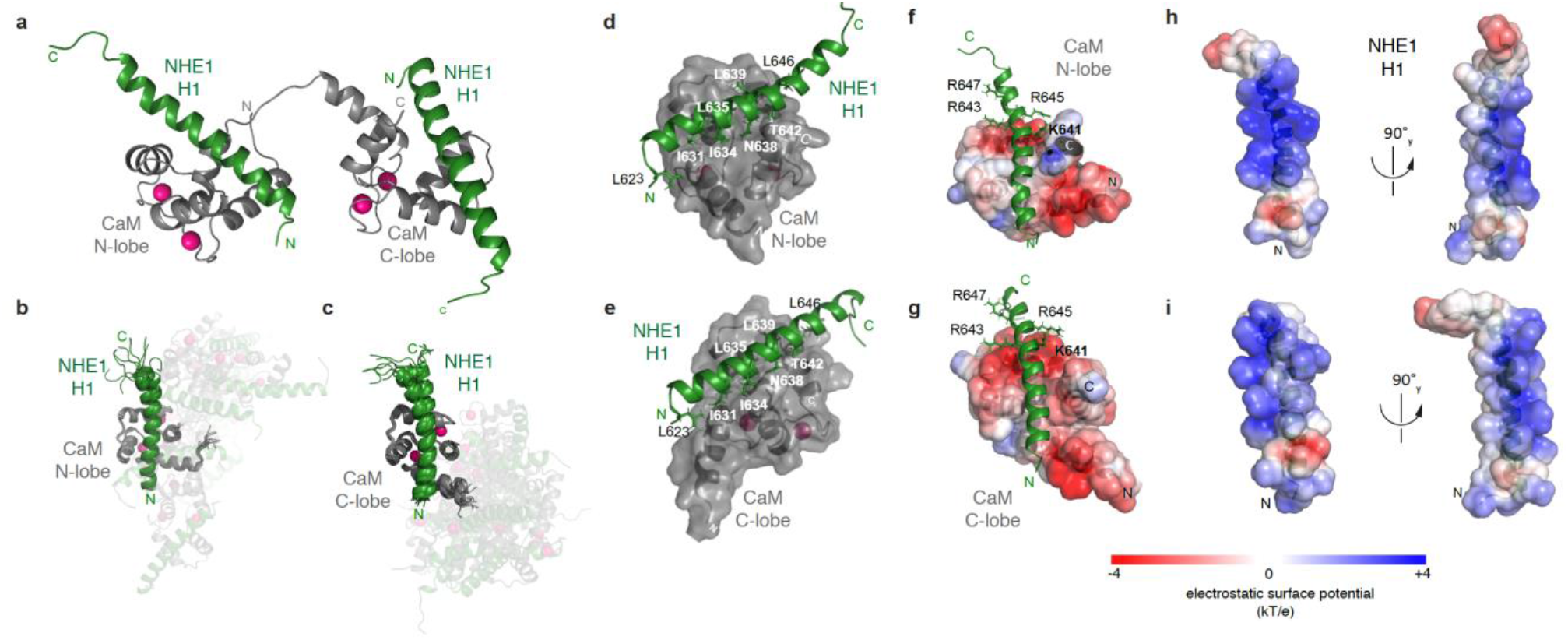
The ternary complex between CaM and two NHE1 H1s. (a) Cartoon of the complex showing the two lobes of CaM in grey and the two NHE1 H1 helices in green. Ca^2+^-ions are shown in magenta. (b,c) Superimposition of the 10 lowest energy structures of the calculated ensemble aligned by (b) the CaM N-lobe with NHE1 H1 bound or (c) the CaM C-lobe with the second NHE1 H1 bound, showing mobility around the CaM linker. (d,e) Hydrophobic residues of NHE1 H1 forming contacts to CaM in the (d) N-lobe and (e) C-lobe, respectively. Electrostatic complementarity between Ca^2+^-CaM and NHE1-H1 in the (f) CaM N-lobe and the (g) CaM C-lobe shown by the surface potential of CaM and (h, i) for the two correspondingly bound H1 helices (pdb accession code 6zbi). **Figure supplement 1:** Comparison of the crystal structure (PDB: 2ygg) and the NMR structure presented here (PDB: 6zbi). **Source data:** PDB coordinates for 6zbi.

### S648 phosphorylation status regulates Ca^2+^-dependent NHE1 activation in cells

The NHE1 C-terminal tail contains several phosphorylation sites some of which are located in the CaM-binding region [26]. S648, which can be phosphorylated by Akt [35,36] is located at the C-terminal distal end of H1. In cardiomyocytes, the S648A mutation increased CaM binding to NHE1 *in vitro*, and S648 phosphorylation by Akt was proposed to inhibit NHE1 by preventing CaM binding [35]. In contrast, in fibroblasts, S648 phosphorylation by Akt was required for NHE1 activation by platelet-derived growth factor [36], indicating that the impact of this residue may be cell-type-and/or context-specific.

Having the structural information described above at hand, we revisited this issue. To determine the effect of Ca^2+^-CaM interaction and S648 phosphorylation on NHE1 localization and function, we generated stable clones of PS120 cells (a mammalian fibroblast cell line which lacks endogenous NHE1, [44]) expressing full-length wild type (WT) hNHE1 and three variants. The 1K3R1D3E (K641D, R643E, R645E, R647E) variation introduces a charge-reversal (CH) of four residues in H1, and will be referred to as H1-CR. A similar charge-reversal variant was reported to exhibit reduced CaM-binding and increased pH_i_ sensitivity, and fibroblasts expressing this variant showed reduced Ca^2+^-induced NHE1 activation [28,29], whereas MDA-MB-231 breast cancer cells showed increased invasion, migration and spheroid growth [45]. The other two NHE1 variants expressed were S648A and S648D, representing the non-phosphorylated and phosphorylated states of S648, respectively. WT NHE1 and all three variants primarily localized to the plasma membrane - the expected pattern for exogenously expressed NHE1 localization in PS120 cells (Fig. 5a) [46–48]. NHE1 protein expression levels were comparable between cell lines, albeit slightly higher in cells expressing the variants compared to WT, as shown by immunoblot analysis (Fig. 5b-c). As expected, endogenous CaM exhibited similar expression levels in all cell lines (Fig. 5b,d), and localized mainly diffusely in the cytosol (Figure 5 – Figure supplement 1).

**Fig 5.**
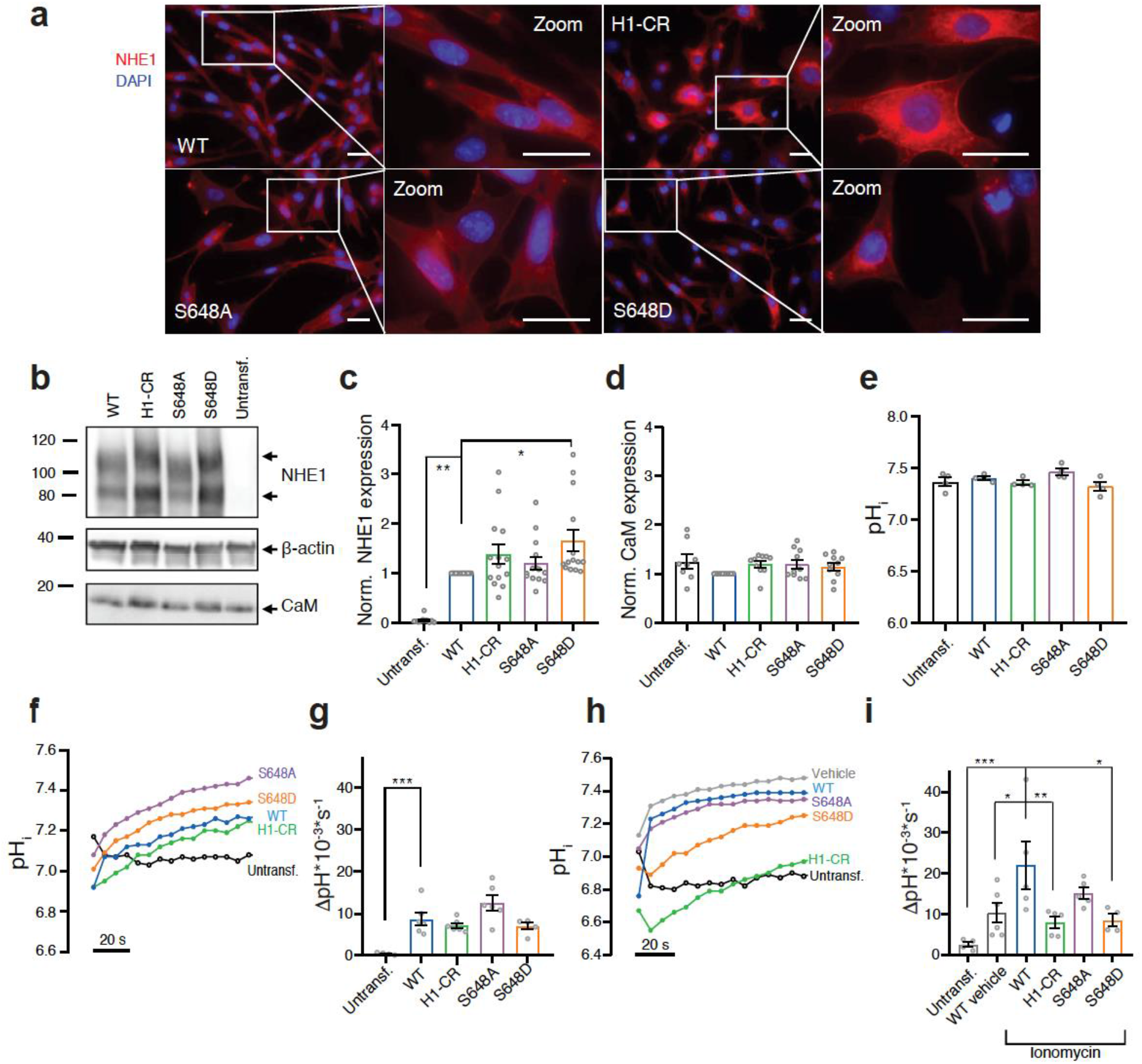
S648 phosphorylation status regulates Ca^2+^-dependent NHE1 activation. (a) Representative IF images of PS120 cells stably expressing WT or variant hNHE1 and stained for NHE1 (red) and DAPI (blue) to evaluate NHE1 expression and localization. Images were taken at 60x magnification on an inverted Olympus Cell Vivo IX83 microscope and areas marked with white squares are magnified. Scale bars represent 20 μm (*n*=*5*). IF images of CaM in all cell lines are shown in Figure supplement 1. (b) Representative Western blot of the four variant PS120 cell lines as well as untransfected PS120 cells blotted for NHE1 and endogenous CaM. For NHE1, the lower band corresponds to the immature protein and the upper band to fully glycosylated NHE1. β-actin was used as loading control (*n*=*8-14*). (c-d) Total NHE1 (*n*=*11-14*) or CaM (*n*=*8-10*) expression was quantified using ImageJ and normalized for each variant to the loading control and then to the expression in the WT NHE1 cell line. (e) Steady-state pH_i_ measured in Ringer solution in the absence of HCO_3_-for each cell line. (*n*=*4*). (f-i) Representative traces of BCECF-AM fluorescence converted to pH_i_ for the four NHE1 variant cell lines and untransfected PS120 cells illustrating the time course of pH_i_ recovery in Ringer in the absence of HCO_3_-after an NH_4_Cl-prepulse-induced acid load without (f-g) or with (h-i) the addition of 5 μM ionomycin or vehicle (EtOH). The recovery rates as ΔpH*10^−3^*s^−1^ of each cell line are depicted in (g) (*n*=*4-6*) and (i) (*n*=*4-6)*. The rate of pH_i_ recovery is expressed as ΔpH/s, because steady state pH_i_ and acidification were similar between cell lines, allowing direct comparison. It should be noted that the variant NHE1 proteins 1K3R1D3E, S648A and S648D harbor a spontaneous F395Y mutation. The expression and pH_i_ recovery rates of two WT NHE1s (two different clones) did not differ from that of F395Y (Figure supplement 2). Data represent mean ± SEM. For c-e, g and i, one-way ANOVA with Dunnett’s post-test comparing all cells to WT was carried out. Exact p-values stated for each group compared to WT were as follows: c) Untransf. p=0.0011, H1-CR p=0.3687, S648A p=0.8909, S648D p=0.0256, d) Untransf. p=0.2897, H1-CR p=0.4609, S648A p=0.4764, S648D p=0.7282, e) Untransf. p=0.8894, H1-CR p=0.7967, S648A p=0.4624, S648D p=0.2813, g) Untransf. p=0.0009, H1-CR p=0.7781, S648A p=0.1097, S648D p=0.7652, i) Untransf. p=0.0007, WT vehicle p=0.0254, H1-CR p=0.0089, S648A p=0.3367, S648D p=0.0195. *, ** and *** denotes p<0.05, p<0.01 and p<0.001, respectively. **Figure supplement 1:** Immunofluorescence staining of endogenous CaM in PS120 cell lines stably expressing WT or variant hNHE1. **Figure supplement 2**: Comparison of expression levels and pH_i_ recovery rates of PS120 cells expressing two WT and F395Y hNHE1.

To determine the effect of CaM binding and S648 phosphorylation status on NHE1 ion transport activity, cells were loaded with the pH sensitive fluorescent probe BCECF-AM and subjected to real-time analysis of pH_i_. The steady-state pH_i_ was similar between all cell lines, ranging from ~7.4-7.5 (Fig. 5e). To directly assess Na^+^/H^+^ exchange activity, we measured the rate of pH_i_ recovery after an NH_4_Cl-prepulse-induced acid load. Since PS120 cells lack endogenous NHE1 activity (Fig. 5f-i, untransf.), pH_i_ recovery in the nominal absence of HCO_3_-reflects the activity of exogenously expressed NHE1 [18]. Following an acid load without further stimulation, recovery rates were not significantly different between cells expressing WT and variant NHE1s (Fig. 5f-g). In the presence of the Ca^2+^-ionophore ionomycin, which causes elevated [Ca^2+^]_i_, concomitant with the acid load, cells expressing WT NHE1 recovered significantly faster than WT vehicle controls (Fig. 5h-i), in agreement with previous reports [29]. Under ionomycin-stimulated conditions, recovery was significantly reduced in cells expressing the NHE1 H1-CR and S648D variants, compared to those expressing WT NHE1 (Fig. 5h-i).

These results show that mutations within the CaM binding region (specifically H1) or mimicking S648 phosphorylation reduce Ca^2+^-stimulated NHE1 activation in PS120 fibroblasts.

### Cellular NHE1:CaM proximity is resistant to H1-CR and S648 NHE1 mutations

To study the interaction between CaM and NHE1 *in situ*, we visualized and quantified their close proximity using proximity ligation assay (PLA), which detects close proximity (≤ 40 nm) of two proteins of interest *in situ* [49]. Previous work has established the feasibility of detecting CaM interactions specifically using PLA and demonstrated that PLA signals between two binding partner proteins can be abrogated by mutation of one protein partner [50,51]. Fig. 6a shows the localization of NHE1:CaM PLA puncta (magenta) and NHE1 (green). Confirming assay specificity, PLA puncta were absent in native PS120 cells not expressing NHE1 (Fig. 6b and Figure 6 – Figure supplement 1). In PS120 cells expressing WT NHE1, PLA puncta, representing ≤ 40 nM proximity of CaM and NHE1, were clearly detected (Fig. 6a-b). The number of PLA puncta per area did not increase upon ionomycin treatment (Fig. 6a-b), suggesting that an increase in [Ca^2+^]_i_ does not detectably increase the number of CaM and NHE1 proteins within close proximity of each other.

**Fig 6.**
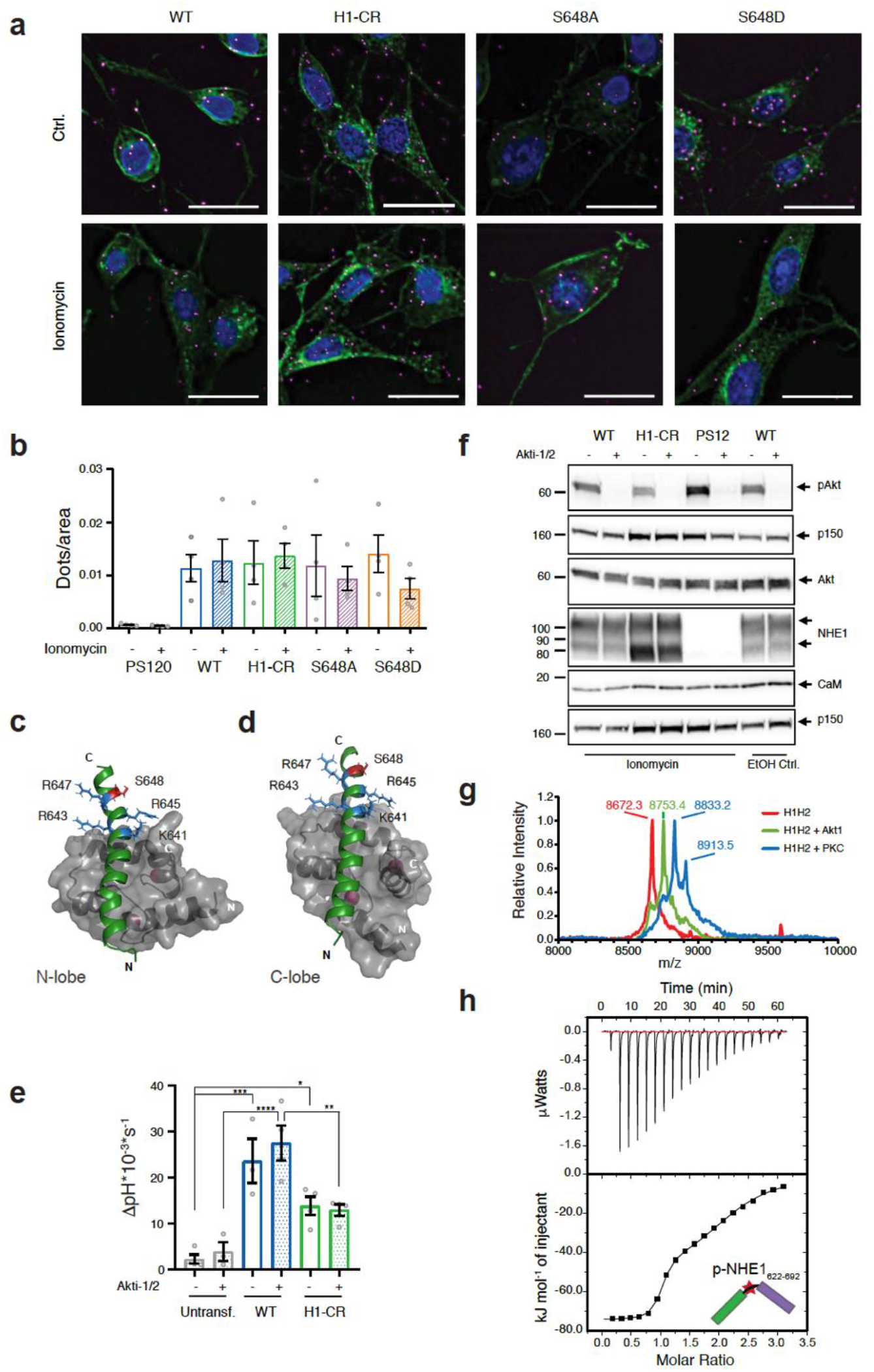
Cellular NHE1:CaM proximity is resistant to H1-CR and S648 NHE1 mutations and Akt activity does not significantly impact NHE1 activity. (a) Proximity ligation assay (PLA) of WT and variant hNHE1 in PS120 cells, with addition of vehicle (EtOH) or 5 μM ionomycin as shown (*n*=*4-7*). Negative controls are shown in Figure supplement 2. PLA signal is magenta, co-staining for NHE1 is green and nuclei (DAPI) are blue. Scale bars represent 20 μm. (b) Quantification of PLA signal in ImageJ after background subtraction and normalization to cell area. Approx. 400 cells were counted per condition per n *(n*=*4)*. Two-way ANOVA with Tukey’s post-test comparing all conditions to one another showed no significant differences between groups. As there are 45 exact p-values for this figure, these are available upon request. (c-d) Structures of the individual lobes of the (2:1) NHE1 H1-CaM complex highlighting the position of the mutated residues and S648 in (c) the N-lobe and (d) the C-lobe. (e) Recovery rates as ΔpH*10^−3^*s^−1^ of each cell line with addition of 10 μM Akti-1/2 or vehicle (DMSO) (*n*=*4*). Two-way ANOVA with Tukey’s post-test comparing cell types to one another within each treatment group. Exact p-values were as follows: e) Ctrl group: Untransf. - WT p=0.0001, Untransf. - H1-CR p=0.0141, WT - H1-CR p=0.0559, Akti-1/2 group: Untransf. - WT p<0.0001, Untransf. - H1-CR p=0.0792, WT - H1-CR p=0.0024, (f) Western blot assessing the inhibition of Akt activity by Akti-1/2, assessed as phosphorylation of Ser473 of Akt (pAkt) compared to total Akt, in PS120 cells expressing WT and variant NHE1 (*n*=*4*). The blot also illustrates endogenous levels of CaM and exogeneous NHE1. p150 was used as a loading control. NHE1 and CaM were detected on the same membrane as Akt. (g) MALDI-TOF MS analyses of NHE1-H1H2 before (red) and after addition of Akt (green) or PKC (blue). (h) Representative binding experiments of Akt-phosphorylated NHE1-H1H2 titrated into CaM using ITC. The upper part shows baseline-corrected raw data from the titrations and the lower part the normalized integrated binding isotherms with the fitted binding curves assuming a two-site binding event. The peptide titrated into CaM is shown in cartoon and the star indicates phosphorylation. The ITC-raw data and the experiments not shown are available in **Source data file** 1.14-1.18. Data represent mean ± SEM. *, **, *** and **** denotes p<0.05, p<0.01, p<0.001 and p<0.0001, respectively. **Figure supplement 1:** Lack of PLA signal in PS120 cells lacking NHE1. **Figure supplement 2:** Identification of phosphorylated residues in NHE1 H1H2 by NMR.

To determine the impact of S648 phosphorylation status on NHE1:CaM interaction in a cellular context, PS120 cells stably expressing WT NHE1, H1-CR, and the two phosphorylation variants, S648A and S648D, were also probed for NHE1:CaM proximity by PLA. In contrast to their inhibitory effect on Ca^2+^-induced NHE1 activation (Fig. 5h-i), none of the three variants abolished NHE1:CaM PLA *in situ* (Fig. 6a-b). We therefore examined the solution NMR structure of the 1:2 (CaM:H1) complex and the positions of the different mutated residues. In the complex, the four residues K641, R643, R645 and R648 are all solvent exposed both when bound to the N-lobe or the C-lobe of CaM, with some hydrophobic contacts from K641 (Fig. 6c,d). Likewise, in either of the lobes, S648 does not form direct contacts with CaM (Fig. 6c,d). While several other mechanisms are possible (see Discussion), this may explain why the H1-CR, S648A and S648D mutations have no apparent effect on NHE1:CaM proximity although both H1-CR and S648D reduced Ca^2+^-induced NHE1 activity.

These results show that despite their effects on Ca^2+^-CaM-induced NHE1 activity, neither H1-CR, S648A nor S648D mutations abrogate the close proximity between NHE1 and CaM in a cellular context.

### Akt activity is not solely responsible for the S648 phosphorylation-dependent NHE1 regulation

In addition to its phosphorylation by Akt [35,36], S648 is phosphorylated or predicted to be phosphorylated by numerous other kinases, including protein kinase C (PKC), protein kinase A, cyclin-dependent kinase, Aurora kinase, and CDC-like kinase-2 [26]. We therefore asked whether Akt was an essential regulator of Ca^2+^-dependent NHE1 activity. A 10 min stimulation with ionomycin had no detectable effect on Akt activation (phosphorylation) in PS120 cells (Fig. 6f). Furthermore, Akt activity was ablated under both control and ionomycin conditions by the Akt inhibitor Akti-1/2 (Fig. 6f), yet Akti-1/2 had no significant effect on ionomycin-stimulated NHE1 activity after an acid load (Fig. 6e). These observations suggest that other kinases than Akt might regulate S648 phosphorylation in the context of an increase in [Ca^2+^]_i_. To investigate this, and to assess how phosphorylation of S648 impacts NHE1:CaM interaction *in vitro*, we phosphorylated the H1H2 peptide using either recombinant Akt or PKCδ in an *in vitro* phosphorylation assay. Phosphorylation of a single residue by Akt was confirmed by mass spectrometry, and S648 was identified by NMR, while PKCδ gave rise to mono-, double- and triple phosphorylated forms of H1H2, identified by NMR at residues T642, S648 and T653 (Fig. 6g and Figure 6 – Figure supplement 2). Thus, H1H2 is a potential target for other kinases. Finally, we used ITC to measure the affinity of monophosphorylated (pS648) H1H2 for CaM. As for the unphosphorylated H1H2 we observed two transitions in the binding isotherm and fitted this to a two-site binding model. The first binding event gave a *K*_*d,1*_ = 5 ± 4 nM with a stoichiometry n_1_ = 0.93 ± 0.02, while the second binding event gave a *K*_*d,1*_ = 2700 ± 600 nM with n_2_ = 1.16 ± 0.06 (Fig. 6h, Table 1), resulting in a 10-fold weakened binding compared to WT.

These results show that under the conditions studied here, inhibition of Akt activity does not alter Ca^2+^-induced cellular WT NHE1 activity. Furthermore, S648 is phosphorylated by several kinases *in vitro*, weakening H1H2 binding to CaM 10-fold. Collectively this indicates that other kinases are important for S648 phosphorylation. Additionally, T642 and T653, present within the binding interface of the H1H2 NHE1 complex, are other phosphorylation sites potentially important for regulating NHE1:CaM interaction.

## Discussion

CaM is highly abundant in cells and its interactome comprises hundreds of proteins, the vast majority of which form 1:1 complexes with CaM [1,2]. Most existing structures of CaM complexes have been solved by crystallography, providing a single, static representation of the interaction. Here, we used solution NMR and reveal that the energy landscape of the CaM complex with the Na^+^/H^+^ exchanger NHE1 - a key regulator of pH_i_ in mammalian cells - is much broader than previously anticipated, populating several different states with different affinities and structures, and modulated by phosphorylation, NHE1:CaM ratio, and [Ca^2+^]_i_ (Fig. 7a-b). *In vitro,* we found that NHE1 and CaM can form either 1:1 or 2:1 (NHE1:CaM) complexes with similar affinities but different structures, suggesting that the complex is highly dynamic and that the relevant state at any time is dependent on the availability of Ca^2+^-CaM and NHE1 in the cell (Fig. 7). In the absence of available Ca^2+^, apo-CaM can associate weakly with NHE1 in a different type of complex (Fig. 7c). A recent study of the interaction between CaM and a truncated, minimal version of eEF-2K showed that the Ca^2+^-free C-lobe bound the truncated eEF-2K in the absence of Ca^2+^. Increasing [Ca^2+^], introduced an avidity effect through additional binding to the Ca^2+^-loaded N-lobe [11]. In a cellular context, where resting [Ca^2+^]_i_ is around 50-80 nM, a low [Ca^2+^-CaM] scenario is more likely than complete absence of Ca^2+^. Under such low Ca^2+^-conditions, our results show that NHE1:CaM interaction can stabilize Ca^2+^-binding to the C-lobe of CaM, resulting in an apo-state N-lobe and a Ca^2+^-loaded C-lobe bound to H1 (Fig. 7d). At higher [Ca^2+^-CaM], corresponding to, e.g., mitogenic stimuli which can increase [Ca^2+^]_i_ to up to around 1 μM, the present study shows that two scenarios are possible: a 1:1 NHE1:CaM complex, with the N-lobe bound to H1 and the C-lobe bound to H2 (Fig. 7e), or a 2:1 NHE1:CaM complex, with the CaM N- and C-lobes both interacting with H1 on two adjacent NHE1 C-tails, potentially stabilizing the dimeric state of NHE1 (Fig. 7f). As CaM is limiting in a cellular context, the likelihood of each of the two latter scenarios is probably modulated by CaM interactions with other proteins at any given time. Thus, although the total cellular [CaM] is 5-6 μM as measured in various cell types, the global free Ca^2+^-CaM concentration in a cell has been estimated to around 50 nM, i.e. ~1% of total [CaM] [52,53]. Both [CaM] and [Ca^2+^]_i_ may, however, reach much higher concentrations locally, e.g. close to Ca^2+^ channels [54], and the precise local Ca^2+^-CaM availability around NHE1 will be dependent on local and global Ca^2+^ signaling and competition with other CaM binding proteins of various affinities.

**Fig 7.**
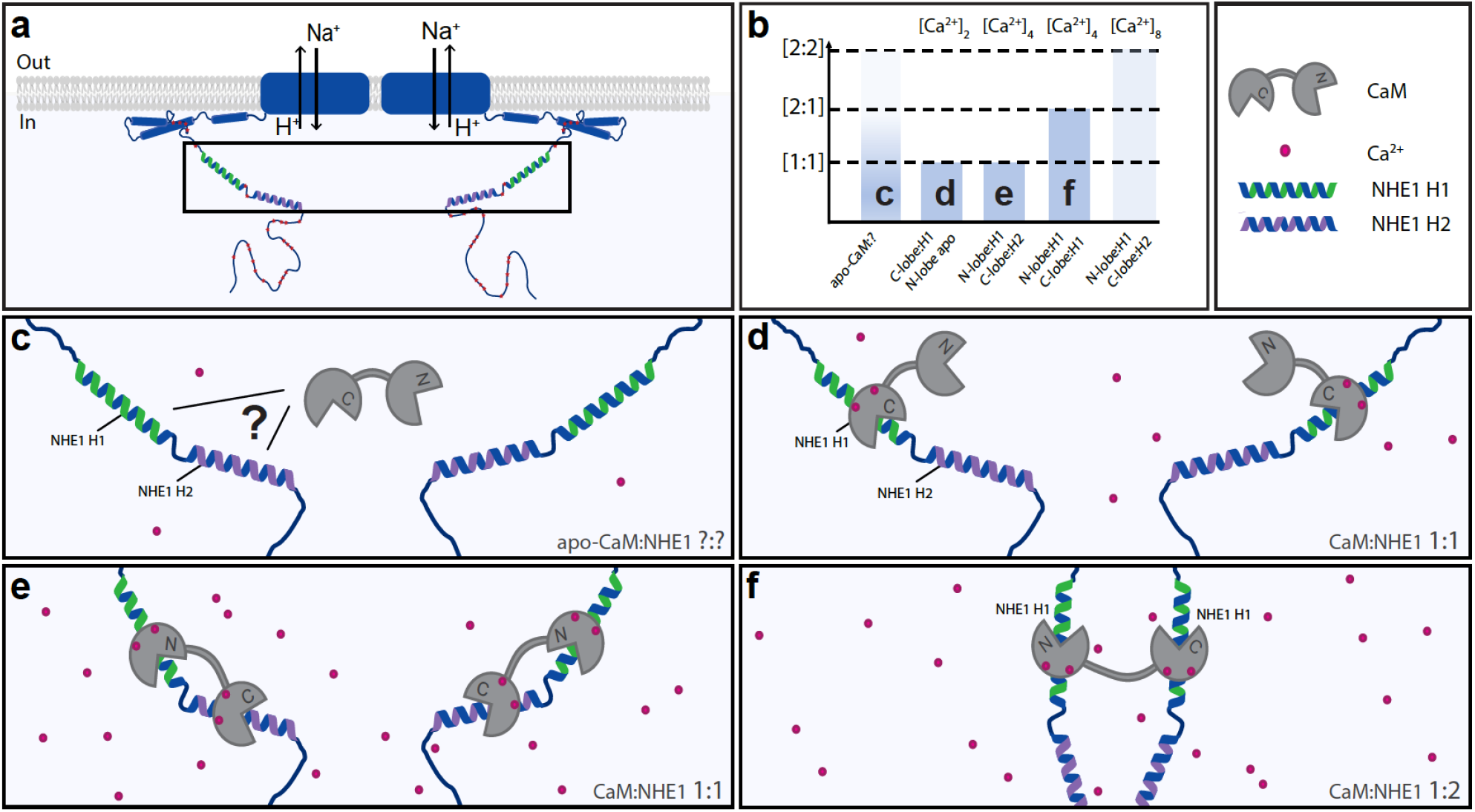
A dynamic view of NHE1:CaM interaction. Multiple possible complexes exist depending on the [NHE1], [CaM], and [Ca^2+^], including ternary complexes, suggesting that CaM may also contribute to NHE1 dimerization. a) Schematic of NHE1 with a zoom on the CaM binding region. The orientation of the monomers (N-terminal to N-terminal) in the NHE1 dimer is arbitrarily chosen. b) Overview of the multiple complexes between CaM and NHE1 and their dependence on Ca^2+^ (top). The indices c), d), e) and f) refer to the states shown in cartoon below. c) Interaction between apo-CaM and NHE1 is seen, but the structure of it is unknown (indicated by question mark). d) At low Ca^2+^-levels, NHE1-H1 is bound to the Ca^2+^-loaded C-lobe of CaM, whereas the N-lobe is in its apo-state. e) at high Ca^2+^-levels, and high NHE1:Ca^2+^-CaM ratio, a 1:1 complex is formed with H1 bound to the N-lobe and H2 bound to the C-lobe of CaM. f) at high Ca^2+^-levels, and low NHE1:Ca^2+^-CaM ratio, a 1:2 complex is formed, where NHE1-H1 is bound to both the N-lobe and the C-lobe of CaM.

CaM-mediated oligomerization has been reported for both soluble and membrane proteins. Examples of the former include the ERα, two of which bind to one Ca^2+^-CaM, one ERα in each lobe, to maximally activate transcription [13,55]. The affinity of ERα for one CaM lobe is almost 100 fold lower than what we see here for NHE1, further supporting a unique role of the 1:1 NH1:CaM complex. For membrane proteins, both CaM-mediated multimerization and apo-CaM binding are well studied for tetrameric, voltage-gated K^+^ channels. Small-conductance Ca^2+^-activated K^+^-(SK) channel monomers form a 1:1 complex with apo-CaM, which binds through the C-lobe and keeps the channel inactive. When [Ca^2+^]_i_ increases, the N-lobe also associates with the channel, triggering dimerization and channel opening [56–58]. Another example is the eag1/Kv10.1 channel, which is inhibited by CaM. Also for this interaction, the stoichiometry is 1:1, and it is proposed that CaM acts as a molecular clamp interacting with domains from neighboring subunits, thereby “twisting” the pore closed [14].

NHE1 proteins are functional homodimers in a manner involving several incompletely understood mechanisms [18]. Based on our findings, it could be speculated that CaM binding contributes to NHE1 C-tail dimerization in a similar way as for ERα, by CaM binding to H1 on two separate NHE1s in a 2:1 (NHE1:CaM) stoichiometry (Fig. 7f), as shown in the ternary complex (Fig. 4). Conceivably, the 1:1 stoichiometry observed for CaM:NHE1_622-692_, with C-lobe:H2 and N-lobe:H1 interaction, could also be possible through a 2:2 complex. This would require each CaM to span two NHE1 C-tails, binding to two different sites on two tails, forming a cross-over structure (not shown). Whether this is feasible in a given cellular context will depend on the structural organization of the C-tail complexed with its binding partners, and on the interaction of the NHE1 lipid interaction domain (LID) with the plasma membrane. Finally, in addition to the direct NHE1:CaM interactions studied here, the spectrum of NHE1:CaM interactions *in vivo* additionally involves indirect interactions, such as the complex between the disordered NHE1 C-tail and the CaM-binding Ser/Thr phosphatase calcineurin [48,59].

Since the discovery that NHE1 is a CaM binding protein [28,29], interaction with CaM has emerged as important for its regulation in response to a wide range of stimuli [32–34]. In most cases, however, the mechanisms involved have remained unaddressed. Initially, release of an autoinhibitory interaction with an allosteric “pH sensor site” on NHE1 by CaM interaction was proposed [29]. However, detailed kinetic analyses have questioned the existence of such a site and indicated that NHE1 activity is regulated by its dimerization state [18,40,42,43]. Our finding that CaM may contribute to stabilization of the NHE1 dimer state would reconcile these models. Finally, given the key role of anionic lipids including phosphatidyl-inositol(4,5)-diphosphate (PI(4,5)P_2_) in regulating NHE1 by interacting with the C-tail [60,61], Ca^2+^-CaM could also regulate NHE1 through electrostatic tuning of the NHE1:PI(4,5)P_2_ interaction. Such mechanisms are reported for several other membrane proteins [62,63] and would be consistent with the folding of the LID domain upon membrane interaction, bringing the CaM binding region in close proximity to the LID and the membrane [64].

The role of phosphorylation of S648 in NHE1 regulation has been a conundrum, with apparently conflicting findings in different cell types [35,36]. Here, we show that phosphorylation of this site which is located at the distant end of H1 reduced NHE1:CaM interaction *in vitro*, yet both the H1-CR variant and the two phosphorylation variants, S648A and S648D retained close (≤ 40 nm) proximity to CaM in cells. This observation mirrors a recent study of CaM binding to smoothelin-like 1 (SMTNL1) protein, where, CaM-SMTNL1 proximity was retained in cells upon mutation, despite loss of interaction *in vitro* [51]. Our results agree with earlier work showing that cellular NHE1:CaM interaction is partially retained in an NHE1 variant functionally similar to our H1-CR variant [28]. Additionally, cellular NHE1:CaM interactions may occur through C-tail interactions with other proteins (e.g. calcineurin) or membrane lipids [48,64], independent of the residues studied here but also giving rise to PLA signals. Finally, while NHE1:CaM proximity in the absence of interaction cannot be excluded, the observed binding of NHE1 to apo-CaM *in vitro*, and the lack of effect of ionomycin on the number of proximity events observed in cells, indicate that a fraction of NHE1:CaM complexes are present at all times.

In congruence with the reduced CaM affinity of the S648-phosphorylated H1H2 *in vitro*, Ca^2+^-induced NHE1 activity in PS120 cells was reduced by the S648D phospho-mimicking variant. Inhibiting the kinase responsible for S648 phosphorylation would therefore be expected to increase Ca^2+^-induced NHE1 activity. In our hands, an increase in [Ca^2+^]_i_ did not activate Akt, one obvious candidate kinase [35,36], and inhibition of Akt had no significant effect on Ca^2+^-mediated NHE1 activity. Furthermore, S648 was phosphorylated *in vitro* by both Akt and PKC, which have overlapping consensus sites, strongly indicating that other kinases than Akt can act *via* S648 to regulate NHE1. Conventional PKCs, which require an increase in [Ca^2+^]_i_ for activation, may be particularly relevant in this context. R643, R645, and R647 which are mutated in NHE1 H1-CR, constitute key residues of the Akt and PKC consensus recognition site [65], which could very well influence phosphorylation in the H1-CR variant. T653 has not been studied in the context of CaM but has been implicated in NHE1 regulation by Rho kinase [66] and by β-Raf [67]. Thus, the mechanisms involved in physiological regulation of NHE1 *via* S648 (and T642 and T653) deserve further studies.

In conclusion, we show that NHE1 and CaM engage in diverse complexes of different NHE1:CaM stoichiometries and structures *in vitro*. The interactions can be dynamically tuned by variations in the concentrations of NHE1, CaM, and Ca^2+^, and by phosphorylation of S648. In cells, Ca^2+^-induced NHE1 activity was reduced by mutation in the first CaM-binding helix, as well as by mimicking S648 phosphorylation, but not by inhibition of Akt. In contrast, NHE1:CaM proximity was not lost by these mutations, likely reflecting further complexity of NHE1:CaM interactions *in vivo*. S648 phosphorylation *in vitro* was facilitated by both Akt and PKC, and Ca^2+^-stimulated NHE1 activity was not affected by Akt inhibition, suggesting that multiple kinases regulate S648 phosphorylation *in vivo*. Finally, we solved the structure of the ternary complex between CaM and two NHE1 peptides, pointing to a possible role of CaM in stabilizing NHE1 dimers *via* binding to two C-terminal regulatory tails. Our results provide novel insight into the dynamics and regulation of NHE1:CaM interaction, and we suggest that an additional NHE1 regulatory layer tuned by Ca^2+^-CaM availability may contribute to NHE1 dimerization. As CaM binding to membrane proteins is widespread, similar regulatory mechanisms and structural diversity may be relevant for tuning oligomerization and function in many other membrane proteins.

## Supplementary material

This article contains online supplementary material.

## Acknowledgements

The authors would like to thank Ruth Hendus-Altenburger for valuable discussion in the early phase of the study, and Johs Dannesboe, Katrine Franklin Mark, Rikke Larsen Agersnap, and Signe A. Sjørup for skilled technical assistance.

## Author contributions

Conceptualization: L.S.F., A.P., B.B.K., and S.F.P. developed the concepts. Methodology, formal analysis and investigation: A.P. and B.B.K. designed, and A.P. and E.S.P. recorded and analyzed the NMR data and assigned the NMR resonances. A.P. did structure calculations, and *in vitro* phosphorylation and mass spectrometry experiments. A.P. and L.S.F. carried out the ITC experiments. J.G.O. did structural analyses, in particular of the crystal structure. L.S.F. and S.F.P. designed all cellular experiments. Cell line generation, western blots, immunofluorescence analysis, and pH_i_ analysis were performed and analyzed by L.S.F. PLA was performed and analyzed by M.S. and L.S.F. Writing, review and editing, and visualization: L.S.F., A.P., E.S.P, B.B.K., and S.F.P. discussed the data, prepared the figures and wrote the manuscript, with comments and inputs from all co-authors. Supervision, project administration and funding acquisition: B.B.K. and S.F.P.

## Funding

This work was supported by a grant from the Danish Research Councils (to B.B.K./S.F.P.: 4181-00344), a Novo Nordisk Foundation SYNERGY grant (B.B.K., S.F.P. #NNF15OC0016670), a Novo Nordisk Foundation Endocrinology grant (S.F.P.), and a Novo Nordisk Scholarship supported L.S.F. We thank Villumfonden for generous support for the NMR instruments. The funding sources were not involved in study design, data collection and interpretation, or the decision to submit the work for publication.

## Conflict of interest

The authors declare that they have no conflicts of interests.

## Materials and Methods

### Antibodies and reagents

Antibody against NHE1 (#sc-136239) was from Santa Cruz Biotechnology, Dallas, TX. Antibodies against CaM (#45689) and (#05-173) were from Abcam, Cambridge, GB, and EMD Millipore, Burlington, MA, respectively. Antibodies against Akt (#9272) and pAkt (Ser473) (#4060) were from Cell Signaling Technology, Danvers, MA. Antibody against p150^Glued^ (#610473) was from BD Transduction Laboratories, Franklin Lakes, NJ. Antibody against β-actin (#A5441 was from Sigma-Aldrich, St. Louis, MO. Anti-mouse (#P0447) and anti-rabbit (#P0448) HRP-conjugated secondary antibodies were from Dako, Glostrup, DK. Anti-mouse-Alexa Fluor 568 (#A10037), anti-rabbit-Alexa Fluor 488 (#A21206) and anti-mouse-Alexa Fluor 488 (#A21202) secondary antibodies for immuno-fluorescence were from Invitrogen, Carlsbad, CA. The Akt inhibitor Akti-1/2 (#124019) was from EMD Millipore.

### Cell lines and cell culture

PS120 cells (a kind gift from Laurent Counillon, University of Nice, France) were grown in DMEM (Life Technologies, Carlsbad, CA) supplemented with 5% (v/v) FBS (Sigma-Aldrich), 100 units and 0.1 mg/mL Pen/Strep (Sigma-Aldrich) and 600 μg/mL G418 (EMD Millipore). The latter only for the transfected cells. Cells were grown at 37oC, 95% humidity, 5% CO_2_ and passaged at a confluence of ~70-100%.

### Stable transfection of PS120 cells with variant NHE1

Transfections were performed with the Lipofectamine 3000 kit (ThermoFisher, Waltham, MA). For each well in a 6-well plate, 125 μL medium with no supplements mixed with 3.75 μL Lipo3000 was added to a mixture of 125 uL medium with no supplements containing 4 μL P3000 reagent and 1 μg plasmid DNA. After 15 min incubation at room temperature (RT), 250 μL of the mixture was added to cells at 40-60% confluency without Pen/Strep and G418 and incubated 5 h. After 5 h, fresh medium containing Pen/Strep was added. The next day, fresh medium containing both Pen/Strep and serum as described was added. At a confluence of ~75-95%, cells were split and grown in medium with the addition of 1900 μg/mL G418 (EMD Millipore). Medium was changed every 2-3 days and cells were split when needed until untransfected control cells had died. To ensure clonality, single colonies of cells arising from a single cell were picked using cell culture cylinders greased with Dow Corning® high-vacuum silicon grease (Sigma-Aldrich). Briefly, cells were thinly seeded in 10 cm petri dishes, washed in PBS and cylinders placed on top of the colonies in question. Then 50 μL 5% (v/v) trypsin (Sigma-Aldrich) was added, cells incubated in the incubator for 5 min and 100-150 μL fresh medium added into the cylinders. Cell suspension from each cylinder was transferred to a well in a 6-well plate and medium added. After clonal selection, the concentration of G418 was decreased to 600 μg/mL.

### Immunofluorescence (IF) analysis

Cells were seeded on 12 mm coverslips in 6-well plates. At a ~70-90% confluence, cells were fixed in 100% MeOH for 15 min on ice and washed (3×5 min) in PBS on ice. Next, the coverslips were placed on parafilm and incubated with 0.5% (v/v) Triton X-100 in TBST (0.01 M Tris/HCl, 0.12 M NaCl, 0.1% Tween 20) for 15 min, blocked in 5% (w/v) BSA in TBST for 30 min and incubated with primary antibodies diluted in 1% (w/v) BSA in TBST for 1.5 h at RT. Coverslips were washed in TBST (3×5 min) and incubated with fluorophore-conjugated secondary antibodies in BSA/TBST for 1.5 h. Coverslips were washed in TBST (4×5 min) including DAPI at dilution 1:1000 in BSA/TBST, mounted with N-propyl-gallate, sealed, and visualized using an inverted Olympus Cell Vivo IX83 with a Yokogawa CSU-W1 confocal scanning unit. Images were processed using ImageJ/Fiji.

### Immunoblotting

Cells were lysed (1% SDS, 10 mM Tris-HCl pH 7.5, 1 mM Na_3_VO_4_, c0mplete protease inhibitor cocktail (Sigma-Aldrich), heated to 95°C), sonicated for 2 × 30 s and centrifuged for 5 min at 20 000 g at 4°C. Total protein amount was determined (DC Protein Assay kit, Bio-Rad, Hercules, CA) and samples equalized with ddH_2_O and NuPAGE LDS 4x Sample Buffer (Invitrogen) with dithiothreitol (DTT) added to final concentrations of 1x (sample buffer) and 125 mM DTT. Samples were run on precast Criterion TGX 10% gels (BioRad) in Tris/Glycine SDS buffer (BioRad) with Benchmark ladder (Invitrogen) and transferred to nitrocellulose membranes (BioRad). Membranes were Ponceau S (Sigma-Aldrich) stained, blocked (5% w/v nonfat dry milk in TBST) and incubated with primary antibodies overnight at 4°C. Then, membranes were washed in TBST for 30 min, incubated with HRP-conjugated secondary antibodies for 1.5-2 h, washed in TBST for 30 min and developed with Pierce ECL Western Blotting Substrate (Thermo Fisher). Band intensities were quantified using ImageJ and normalized to the respective controls.

### Measurements of pH_i_

Measurements of pH_i_ were performed using the ammonium pre-pulse technique. Briefly, 8*10^4^-12*10^4^ PS120 variant cells were seeded in 24-well plates 24 h prior to the experiment, then loaded with 2’,7’-bis-(2-carboxyethyl)-5-(and-6)-carboxyfluorescein acetoxymethylester (BCECF-AM, 1.6 μM, Life Technologies) for 30 min at 37°C. Cells were washed in Ringer solution (in mM, 130 NaCl, 3 KCl, 20 Hepes, 1 MgCl_2_, 0.5 CaCl_2_, 10 NaOH pH 7.4) twice, then bathed in Ringer solution again and placed in a FluoStar Optima plate reader thermostatted at 37°C. Emission was measured at 520 nm and excitation at 485 nm. After baseline measurement (10 min) in Ringer solution, acidification was induced by washout of either 20 mM NH_4_Cl after 5 min exposure (Fig. 5f) or 5 mM after 25 min exposure (Fig. 5h). Na^+^ free solution (in mM, 135 NMDGCl, 3 KCl, 20 Hepes, 1 MgCl_2_, 0.5 CaCl_2_, pH 7.4) was applied (1.5 min) then the recovery was measured in Ringer solution for 10 min (Fig. 5f). For the ionomycin experiment, 5 μM ionomycin (Sigma-Aldrich) or 96% EtOH (final dilution 0.48% v/v) was added to the Na^+^ free and normal Ringer solutions during recovery and recovery was measured for 5 min (Fig. 5h). For the Akti-1/2 experiment, 10 μM Akti-1/2 or DMSO (final dilution 0.11 % v/v) was added to the cells while incubating for 30 min with BCECF-AM as well as to the Ringer and NH_4_Cl solutions and 5 μM ionomycin or 96% EtOH (final dilution 0.48% v/v) was added to the Na^+^ free and the Ringer solutions during recovery which was measured for 5 min (Fig. 6e). Calibration was performed using the high-K^+^/nigericin technique (high K+ Ringer: in mM, 140 KCl, 10 Hepes, 1 MgCl_2_, 0.5 CaCl_2_, 50 μM nigericin, pH 7.4) and an 8-point calibration curve obtained by fitting to the expression 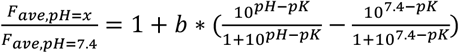 where b and *pK* are constants used for the conversion from fluorescence to pH 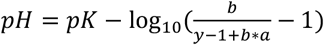 where 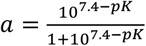 [68,69]. Maximal acidification did not differ significantly between cell lines. Recovery rate was determined by fitting data to the expression IF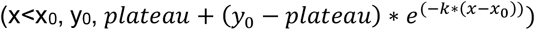 and calculating the derivative at 6 s after addition of Ringer recovery solution. Traces with obviously unacceptable signal to noise ratio and traces showing recovery during the Na+ free step were excluded to ensure that only full initial NHE1 activity was recorded. Recovery rate for PS120 untransfected controls as well as a few traces that were not properly fitted using the expression above (H1-CR in n=2 and n=7) was calculated as the slope of the initial approximately linear part of the curve.

### Proximity ligation assay (PLA)

Cells were seeded on coverslips and treated for 10 min with either 96% EtOH (final dilution 0.48% v/v) in Ringer solution or 5 μM ionomycin in Ringer solution at a confluence of ~50-70%. Then cells were fixed and permeabilized as described for IF analysis. The assay was carried out using Duolink® PLA kit (Sigma-Aldrich) following manufacturer’s protocol. Briefly, blocking solution was added for 25 min at RT, coverslips were then incubated with primary antibodies for 1h at 37°C and washed 3 × 5 min in TBST in a 6-well plate with gentle shaking. PLA probes (PLUS and MINUS) were then added and incubated for 1h at 37°C followed by washing as described. Ligase solution was added and incubated for 30 min at 37°C followed by washing. Amplification solution containing polymerase was then added and incubated for 100 min at 37°C followed by washing protected from light. In some experiments, green anti-mouse fluorophore-conjugated secondary antibody was applied over night at 4°C to stain NHE1. The next day coverslips were washed and DAPI diluted 1:1000 in ddH_2_O was added for 5 min at RT followed by washing. Coverslips were then mounted with N-propyl-gallate, sealed, and visualized using an inverted Olympus Cell Vivo IX83 with a Yokogawa CSU-W1 confocal scanning unit or, for z-stacks, an Olympus BX63. Coverslip identity was blinded for the observer. Confocal images were captured in a single plane across a grid of each coverslip. Images were quantified with ImageJ using the Analyze Particles function on binary images with background subtraction. Cells on coverslips used for quantification (Fig. 6d) were not stained for NHE1. Approximately 400 cells were counted per condition per experiment. Cell sizes were measured manually using merged z-stacks of coverslips stained for NHE1 (Fig. 6a).

### Protein expression and purification

A modified pET24a-vector coding for His-SUMO-tagged human NHE1 A622-D657 (H1), NHE1 R651-D692 (H2) or NHE1 A622-D692 (H1H2) was transformed into *E. coli* BL21 Rosetta 2 (DE3)pLysS cells and grown overnight (o/n) at 37°C in 10 mL LB media with 50 μg/mL kanamycin (kan) and 35 μg/mL chloramphenicol (cam). For expression of unlabeled NHE1 peptides, the cultures were added to 1 L LB media with 50 μg/mL kan and 35 μg/mL cam and grown at 37°C, 185 rpm to an OD_600_ of 0.6-0.8 and induced with 0.5 mM IPTG and harvested after 3 h by centrifugation at 4,700 g for 10 min at 4°C. For expression of stable isotope labelled NHE1 peptides, the o/n culture was added to 1 L M9 minimal media (22 mM KH_2_PO_4_, 42 mM Na_2_HPO_4_·2H_2_O, 17 mM NaCl, 1 mM MgSO_4_, added 1:1000 of M2 trace element solution, 20 mM glucose (if required: ^13^C-labelled), 19 mM NH_4_Cl (if required: ^15^N-labelled) with 50 μg/mL kan and 35 μg/mL cam and grown similar to LB media cultures. After resuspension in 25 mL 50 mM Tris-HCl, pH 7.4 and centrifugation at 5.000 x g for 15 min at 4°C the pellet was dissolved in in 25 mL buffer NA (50 mM Tris-HCl, 10 mM imidazole, 150 mM NaCl, pH 8.0) supplemented with one tablet ethylenediaminetetraacetic acid (EDTA)-free protease inhibitor (Roche). The cell solution on ice was sonicated 3 × 1 min at 90% amplitude (0.5 s cycles) using a UP200S Ultrasonic Processor (Hielscher), with 1 min breaks in between to cool down the sample. Subsequently, phenylmethylsulfonyl fluoride (PMSF) was added to the sample at a final concentration of 1 mM, and centrifugated at 20.000 x g for 45 min at 4°C and the pellet discarded. A total of 5 mL Ni^2+^ Sepharose 6 Fast Flow (GE Healthcare) was transferred to a 20 mL gravity flow column. The lysate supernatant was added to the column and incubated for 1 h at RT under gentle shaking. The column was washed with 50 mL buffer NB (50 mM Tris-HCl, 10 mM imidazole, 1 M NaCl, pH 8.0), followed by a wash with 50 mL buffer NA. The protein was eluted with 10 mL buffer NC (50 mM Tris-HCl, 300 mM imidazole, 150 mM NaCl) and subsequently β-mercaptoethanol was added to a final concentration of 3 mM together with ~100 μg Ubiquitin-like-specific protease (ULP1) and incubated for 1 h at RT. After cleavage the NHE1 peptides were further purified by means of reversed-phase chromatography. A 17.35 mL Zorbax 300Å StableBond C18 column was utilized using an Äkta Purifier 10 HPLC system. TFA was added to the samples containing NHE1 peptides to a final concentration of 0.1% and centrifugated at 20.000 x g for 20 min at RT to remove insoluble aggregates prior to application. The column was equilibrated with 100% buffer RA (0.1% trifluoroacetic acid (TFA) (v/v) in MQ water) and a total volume of 5 mL was applied over two injections. A step gradient from 0% to 100% buffer RB (0.1% TFA (v/v), 70% acetonitrile (v/v) in MQ water) was set over 10 CV with a flow of 3.5 mL/min. A_280_ was measured and fractions of 0.5 to 2.5 mL were collected and analyzed by SDS-PAGE. Fractions containing NHE1 peptides were pooled, flash-frozen, lyophilized o/n and stored at −20°C.

A pET5-vector coding for full length bovine CaM (1-149, 100% protein sequence identity to human CaM), was transformed into *E. coli* BL21 CodonPlus (DE3)-RP cells and grown o/n at 37°C in 10 mL LB media with 100 μg/mL ampicillin (amp) and 35 μg/mL chloramphenicol (cam). 1L of culture medium (LB or M9) was inoculated with 10mL of the overnight culture and expression was induced with 0.5 mM IPTG at an OD ~0.6. After induction the cells were grown for 4h at 37°C and harvested by centrifugation at 6000 g for 10 min. The cell pellet was suspended in 30 mL loading buffer (20 mM Tris-HCl, 500 mM NaCl, 10 mM CaCl_2_, pH 7.5) supplemented with one pellet EDTA-free protease inhibitor (Roche), and sonicated on ice for 5 × 30 s at 80% amplitude (0.5 s cycles) using a UP200S Ultrasonic Processor (Hielscher), with 30 s breaks in between to cool down the sample. Subsequently, the lysate was centrifuged at 20.000 x g for 45 min at 4°C and the pellet discarded. The lysate supernatant was filtered using a 0.45 μm syringe filter and loaded onto a 5mL phenyl Sepharose column. The column was subsequently washed with 40 mL loading buffer before eluting the protein in fractions of 2 mL using a 10 mL gradient from 0-100% elution buffer and additional 10 mL 100% elution buffer (20 mM Tris-HCl, 500 mM NaCl, 50 mM EDTA, pH 7.5). A_280_ was measured and fractions were collected and analyzed by SDS-PAGE. The samples containing CaM were concentrated to ~1 mM and the buffer was exchanged to the Gelfitration buffer (20 mM Tris, 100 mM KCl, optional 2 mM CaCl_2_ depending if the Ca bound or unbound form is desired, pH 7.5), using an Amicon centrifugal filter (10 kDa cutoff) and applied to a HiLoad16/600 Superdex75 column. A_280_ was measured and fractions were collected and analyzed by SDS-PAGE. Fractions containing CaM were pooled and stored at −20°C.

### NMR spectroscopy

All NMR spectra were recorded on Bruker Avance III 600 MHz or 750 MHz spectrometers equipped with TCI cryo-probes. If not specified otherwise, the sample conditions were 20 mM Tris, 100 mM KCl, 2 mM CaCl_2_, 5% D_2_O, 125 μM DSS (adjusted to pH 7.5 with 1 M HCl) at a temperature of 37°C. All spectra were referenced to DSS in the ^1^H direct dimension and indirectly in the ^15^N and ^13^C dimensions using the gyromagnetic ratios. All spectra were zero-filled, apodized using a cosine bell window function in all dimensions, Fourier transformed, and phase corrected manually using either TopSpin®3.6, NmrPipe [70] or qMDD [71], if spectra were recorded using non-linear-sampling (NLS). All spectra were analyzed and assigned manually using CCPNmr Analysis 2.4.2 [72]. For titrations experiments, series of^1^H-^15^N HSQC NMR spectra of either protein (^15^N-labelled) were recorded at 37°C in the absence and with increasing concentrations of the other up to 1:2.5 molar ratios, keeping the detected protein concentration constant.

### NMR resonance assignment

Backbone and sidechain resonances of free CaM (**Sample A**: 1.2 mM ^13^C,^15^N labelled CaM), free H1 (**Sample B:** 0.5 mM ^13^C,^15^N labelled H1, recorded at 5°C), free H2 (**Sample C:** 0.5 mM ^13^C,^15^N H2, recorded at 5°C), as well as of the CaM:H1 complex (**Sample D:** 0.5 mM ^13^C,^15^N labelled CaM + 1.15 mM unlabelled H1; **Sample E:** 1 mM ^13^C,^15^N labelled H1 + 0.5 mM unlabelled CaM) were assigned using sets of triple resonance spectra HNCO, HN(CA)CO, HNCACB, NH(CO)CACB, HcCH-TOCSY, hCCH-TOCSY, ^15^N-NOESY-HSQC, ^13^C-NOESY-HSQC.

### 2D NMR line shape analysis

The ^1^H-^15^N-HSQC spectra were processed using an exponential weighting function in both dimensions (4 and 8 Hz line broadening in F1 and F2, respectively) and analyzed using the 2D lineshape analysis tool TITAN for matlab [73] with a two-state binding model for the titration of CaM with H2. For the titration of CaM with H1 a model with two independent sites (four states) was used for fitting. To reduce the number of variables the K_d_’s as determined from ITC were kept as constants in the fitting process and the assumption that the relaxation rates R_2_(^1^H) and R_2_(^15^N) of a residue in one lobe are not affected by peptide binding to the other lobe. ‮ In both cases, >20 isolated peaks were analyzed, and the errors were determined by a bootstrap analysis and are given by the standard deviation from the mean from 100 replicas.

### Structure calculations and refinements

Intra- and intermolecular distance restraints were obtained from Sample D and E using a set of heteronuclear 3D NOESY experiments (^15^N-NOESY-HSQC, ^13^C-NOESY-HSQC as well as ^12^C/^14^N filtered versions yielding exclusively intermolecular NOEs. The mixing time in all NOESY experiments was 120 ms. Backbone angular restraints were calculated form chemical shifts using Talos N [74]. Initial NOE assignments and structure calculations for the CaM:H1 complex were done iteratively using Cyana 3.98.5 [75]. Four Ca^2+^ ions were included and the geometry of calcium coordination was fixed based on distance and angular restraints from crystal structures (pdb accession code: 1CLL; [76]). The structures were further refined with an implicit water model (eefxPot [77]) using XPLOR-NIH v2.44 [78].

### Isothermal titration calorimetry (ITC)

All ITC data was recorded on a microcal instrument ITC200 (Malvern). Data were recorded at 25°C with a stirring speed of 307 rpm and a reference power of 10 μCal/s. The buffer composition was 20 mM Tris (adjusted to pH 7.5 with 1 M HCl), 100 mM KCl, 2 mM CaCl_2_ and all samples were degassed freshly before usage. For the high affinity interactions (H1, H1H2, and H1H2 pS648), CaM was at an initial concentration of 10 μM in the sample cell and the peptides were added from a stock concentration of 150 μM in the syringe. The titration of H2 into CaM was conducted at 50 μM CaM in the cell and 670 μM H2 in the syringe. In this case the heat of dilution was determined from a control experiment, where H2 was injected into a buffer solution and this dataset was subsequently used for baseline correction. Data from the ITC experiments were analyzed using the Origin 7 software package (MicroCalTM). The datasets were fitted to models for a single binding site (H1, H2), or two independent binding sites (H1, H1H2, H1H2 pS648). The extracted thermodynamic parameters (Table 1) are the mean and standard deviation of three independent experiments and one representative experiment is shown in Fig. 1c-e.

### Helix propensity

Helix propensity was predicted using Agadir (http://agadir.crg.es/) using standard settings [79].

### *In vitro* phosphorylation and mass spectrometry

*In vitro* phosphorylation assays were carried out with 100 μM unlabelled or ^13^C,^15^N labelled H1H2 in 50 μL or 200 μL phosphorylation buffer (25 mM 3-(N-morpholino)propanesulfonic acid (MOPS), 25 mM MgCl_2_, 5 mM ethylene glycol-bis(β-aminoethyl ether)-N,N,N′,N′-tetraacetic acid (EGTA), 2 mM EDTA, 10 mM adenosine triphosphate (ATP), 5 mM dithiothreitol (DTT), pH = 7.2), respectively. To this, 0.3 μg of active, GST-tagged, human Akt1 (Sigma-Aldrich) or 0.3 μg active, GST-tagged, human PKC δ (Biaffin GmbH) was added and incubated at 37°C for up to 120 h. The unlabelled samples were analyzed by Matrix assisted laser desorption/ionization time of flight mass spectrometry (MALDI-TOF MS) on a Bruker autoflex III smartbeam spectrometer with α-cyano-4-hydroxycinnamic acid (HCCA) as matrix after 24 h. The ^13^C,^15^N labelled samples were analyzed by NMR spectroscopy after 120 h and partially assigned as described above.

### Statistical analysis

Data are shown as representative images or as means with standard error of means (SEM) error bars as indicated. Statistical analysis was carried out with software Graph Pad Prism 8. Statistical significance for experiments with one variable (e.g. cell type/group) was assessed using one-way analysis of variance (ANOVA) followed by Dunnett’s or Tukey’s post-test where appropriate (applied for pH_i_ and protein expression data in Fig. 5). Statistical significance for experiments with two variables (e.g. cell type and treatment) was assessed using unmatched two-way ANOVA followed by Tukey’s post-test (applied for pH_i_ and PLA data in Fig. 6). The comparisons carried out and the exact p-values from each test are indicated in the figure legends except for Fig. 6b where the 45 exact p-values are available upon request. *, **, *** and **** denote p < 0.05, 0.01, 0.001 and 0.0001, respectively, unless otherwise specified.

## Additional files

### Figure supplements for main figures

**Figure 2 – figure supplement 1.**
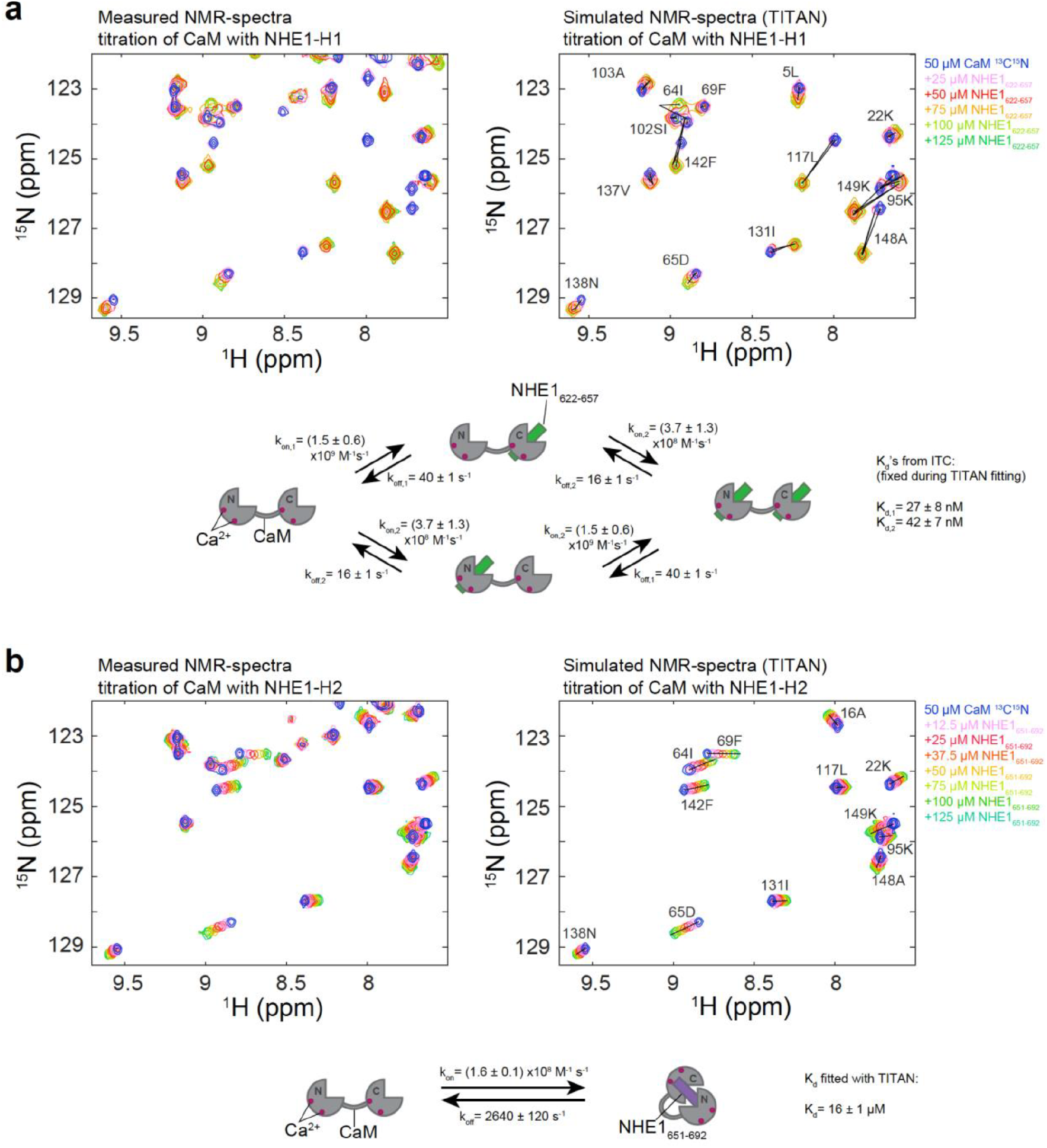
TITAN 2D-NMR lineshape analysis for extraction of binding kinetics. (a) Side-by-side comparison of measured (left) and simulated (right) ^1^ H,^15^N HSQC spectra of 50 μM CaM (^13^C, ^15^ N) titrated with up to 125 μM unlabeled NHE1_622-657_. The simulated spectrum was obtained by fitting the data to a four-state model as depicted in the cartoon below with the parameters for *k*_*off*_ obtained from the fit and the *K*_*d*_s taken from the ITC analysis. (b) Side-by-side comparison of measured (left) and simulated (right) ^1^H,^15^N HSQC spectra of 50 μM CaM (^13^C, ^15^N) titrated with up to 125 μM unlabeled NHE1_651-692_. The simulated spectrum was obtained by fitting the data to a two-state model as depicted in the cartoon below with the parameters *k*_*off*_ and *K*_*d*_ obtained from the fit. The black lines in the simulated spectra connect the fitted signal positions of the four or two states in (a) and (b) respectively.

**Figure 2 – Figure supplement 2.**
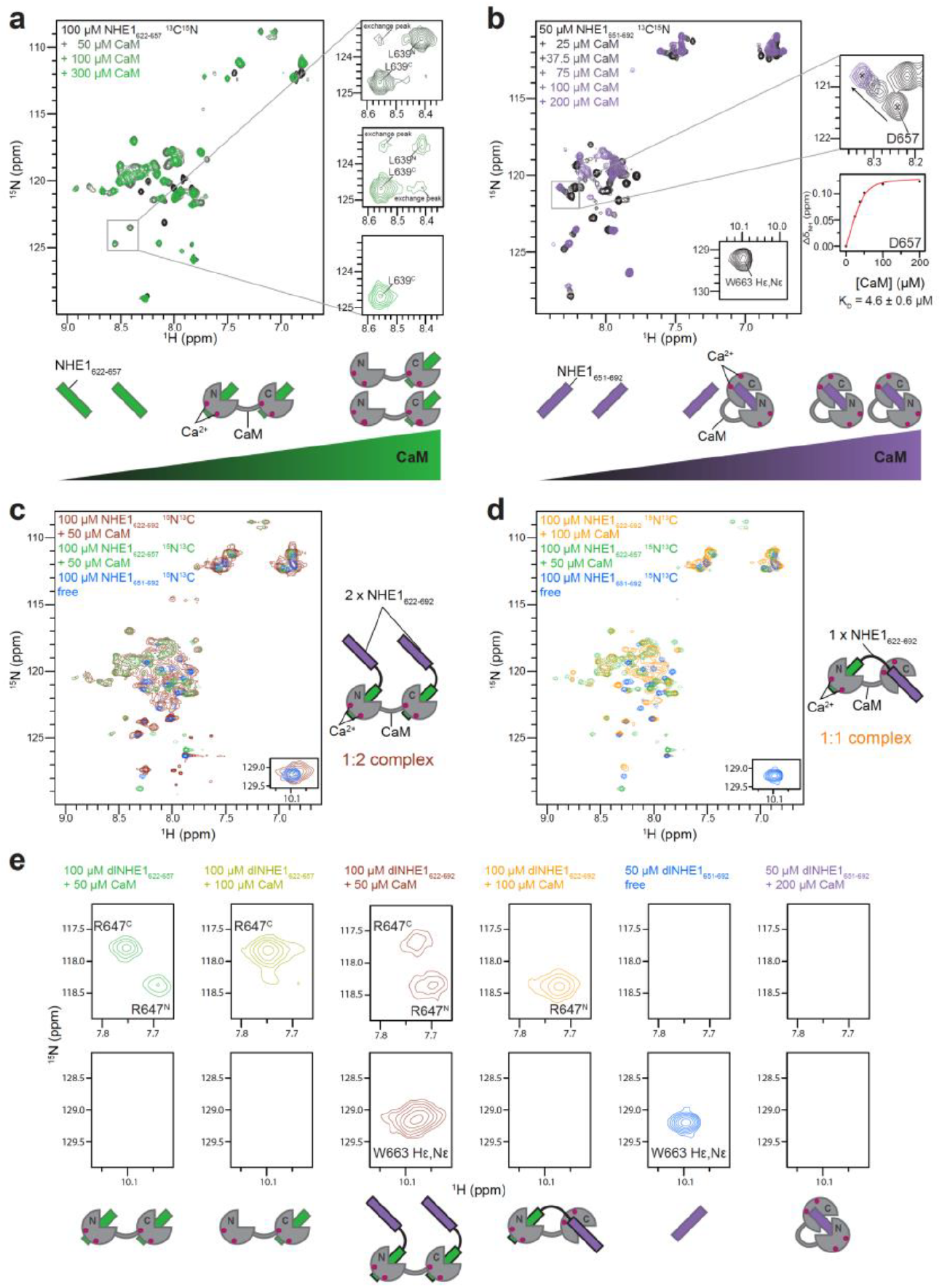
Titration of ^15^N labeled H1, H2 and H1H2 with unlabeled CaM. (a) ^1^H,^15^N HSQC spectra of 100 μM ^15^N-NHE1_622-657_ titrated with up to 300 μM (black to green) unlabeled CaM. The NMR signals of L639 at three different NHE1_622-657_:CaM ratios; 1:1 (top), 1:2 (middle) and 1:3 (bottom) are shown to the right. N and C indicate if the highlighted residue is bound to the N- or C-lobe of CaM. Weak exchange peaks indicate intermediate/slow exchange (~0.1 s^−1^) of NHE1_622-657_ between the two CaM lobes during the experiment. The cartoon representation depicts the most prevalent NHE1_622-657_:CaM states as a function of increasing [CaM]. (b) ^1^H,^15^N HSQC spectra of 50 μM ^15^N-NHE1_651-692_ titrated with up to 200 μM (black to purple) with unlabeled CaM. **NMR**shifts of D657 shown to the right with a fit of the CSPs as a function of CaM concentration, yielding a *K*_*D*_ of 4.6 ± 0.6 **μM.**The cartoon representation depicts the most prevalent NHE1_651-692_:CaM states as a function of increasing [CaM]. (c) ^1^H,^15^N HSQC spectra of 100 μM ^15^N- NHE1_622-692_ in the presence of 50 μM unlabeled CaM. The signals are severely broadened and overlapping but the resonances closely resemble signals from the spectrum of H1 bound to both lobes of CaM (green) or free H2 (blue), confirming the 1:2 (CaM:H1) complex as depicted in the cartoon to the right. (d) ^1^H,^15^N HSQC spectra of 100 μM ^15^N-NHE1_622-692_ in the presence of 100 μM unlabeled CaM. The signals closely resemble resonances from the spectrum of H1 bound to the N-lobe of CaM (green) and few signals resemble resonances from the far C-terminal region of free H2 (blue), confirming the 1 :1 (CaM:H1) complex as depicted in the cartoon to the right. (e) Zooms of the 1H,15N HSQC spectra highlighting a representative resonance of H1 (R647, top row) and H2 (HεNε W663, bottom row). The cartoons depict the most prevalent NHE1:CaM state at the given conditions. di denotes double-labeled (^13^C,^15^N).

**Figure 2 – Figure supplement 3.**
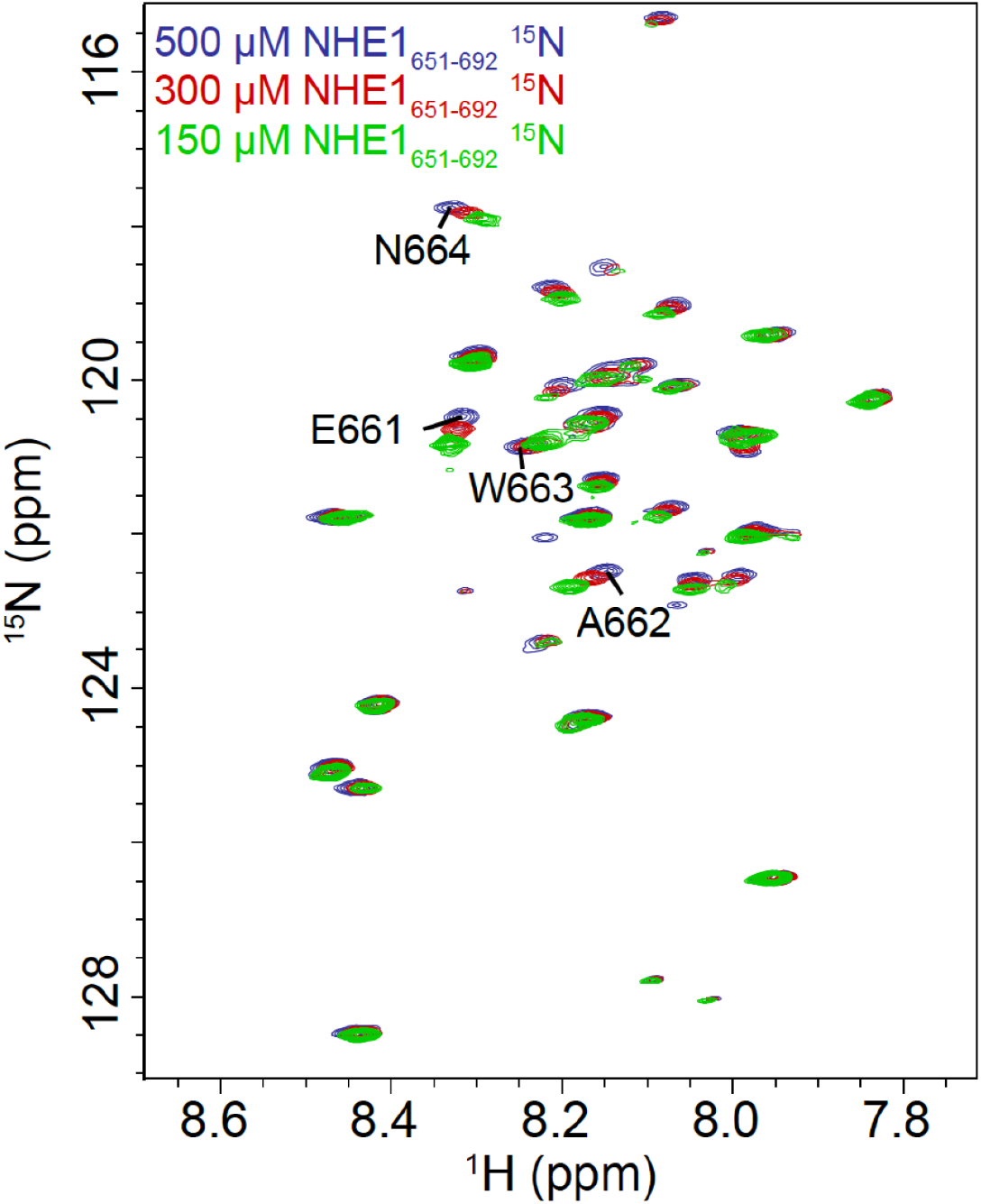
Concentration dependence of the NMR chemical shift of NHE1 H2. ^1^ H, ^15^N HSQC spectra of NHE1 H2 at different concentrations show concentration dependent changes in chemical shifts indicating a weak self-association of the peptide at high concentrations. The most affected resonances are highlighted and correspond to the stretch of residues E661-N664 of NHE1.

**Figure 4 – Figure supplement 1.**
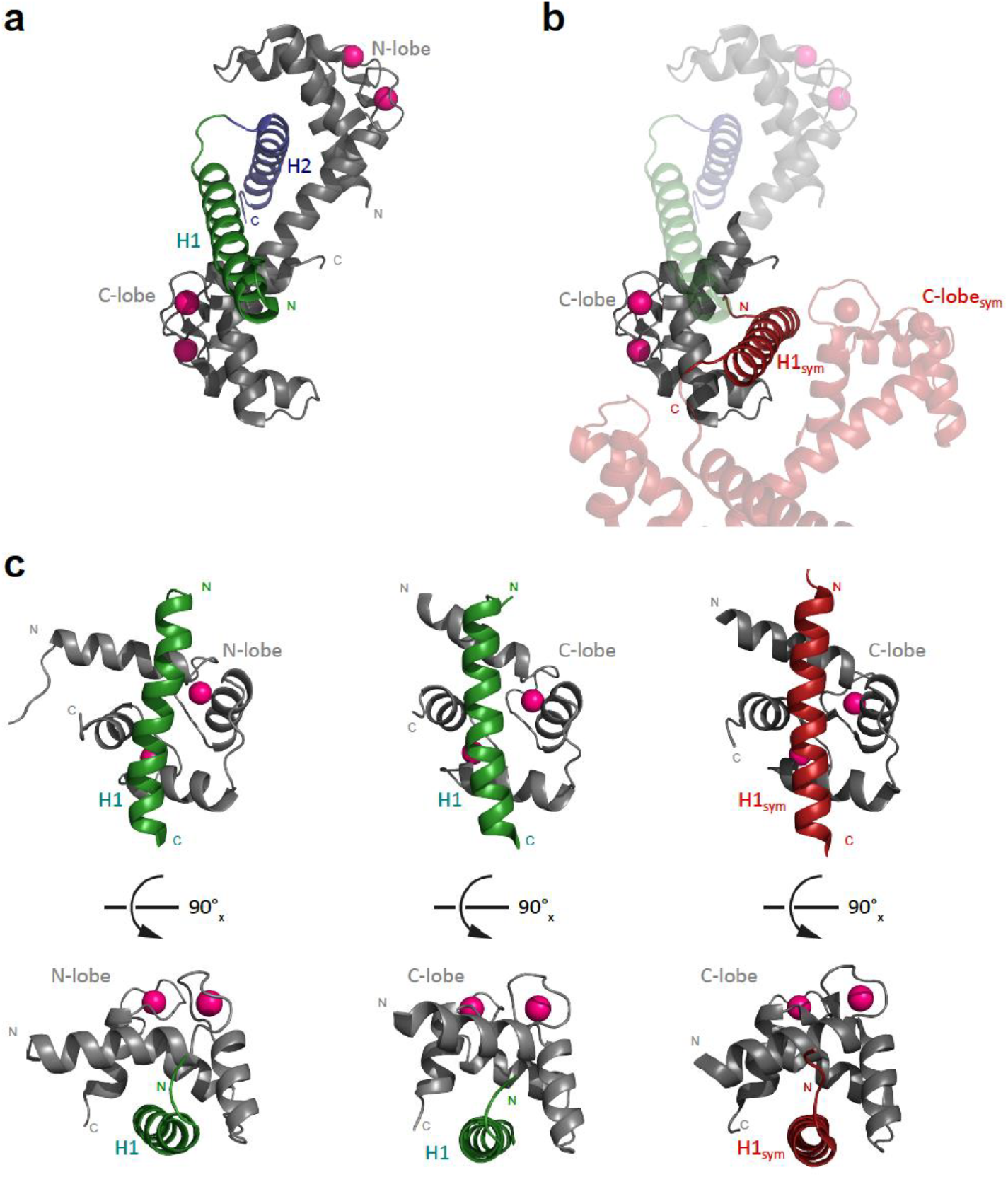
Comparison of the crystal structure and the NMR structure of NHE1:CaM complexes. (a) Crystal structure of the 1:1 NHE1:CaM complex (PDB-code: 2ygg) with the N-lobe bound to H2 and the backside of the C-lobe bound to H1 (PBD code: 6zbi). (b) Same as in (a) with a symmetry-related molecule of the crystal lattice included (red), showing that the hydrophobic cleft of the CaM C-lobe is occupied with H1 of this symmetry-related complex (H1_sym_). (c) Side-by-side comparison of the structures of CaM N-lobe bound to H1 (NMR-structure, left), CaM C-lobe bound to H1 (NMR-structure, middle) and CaM C-lobe bound to H1_sym_ (crystal structure, right), showing the same binding mode in all three cases. The C^α^-RMSDs between the different structures are 2.3 Å (N-lobe+H1_NMR_:C-lobe+H1_NMR_, left:middle), 1.8 Å (N-lobe+H1_NMR_:C-lobe+H1_sym,X-ray_, left/right) and 1.1 Å (C-lobe+H1_NMR_:C-lobe+H1_sym,X-ray_ middle-right), respectively.

**Figure 5 – Figure supplement 1.**
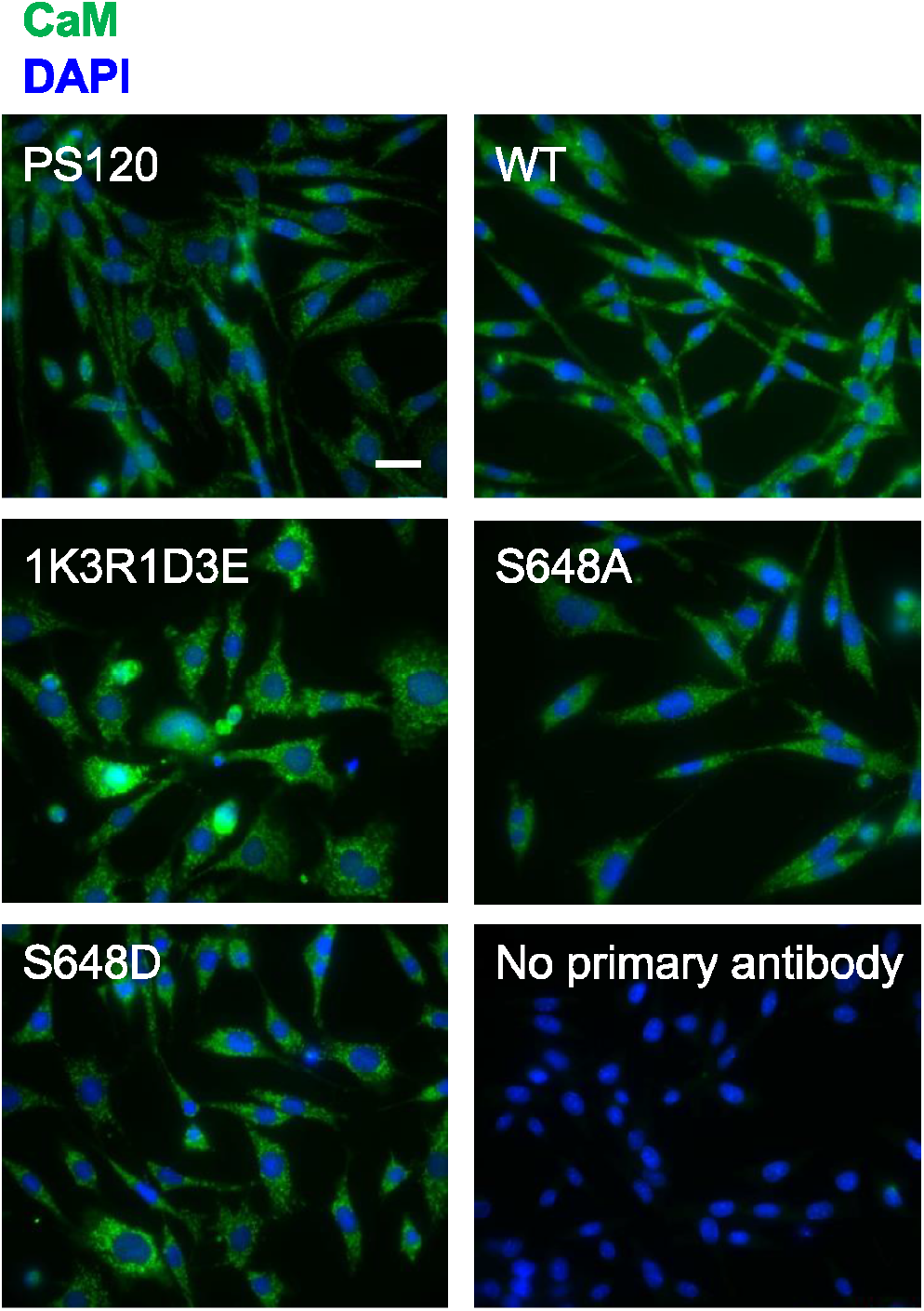
lmmunofluorescence staining of endogenous CaM in PS120 cell lines stably expressing WT or variant hNHE1. Representative IF images of PS120 cells stably expressing WT or variant hNHE1 and stained for CaM (green) and DAPI (blue) to evaluate endogenous CaM expression and localization. Images were taken at 60x magnification on an Olympus Cell Vivo IX83 microscope and are from the same cell passages as the pictures shown in Fig. 5a (main text). Scale bars represent 20 μm *(n*=*5)*.

**Figure 5 – Figure supplement 2.**
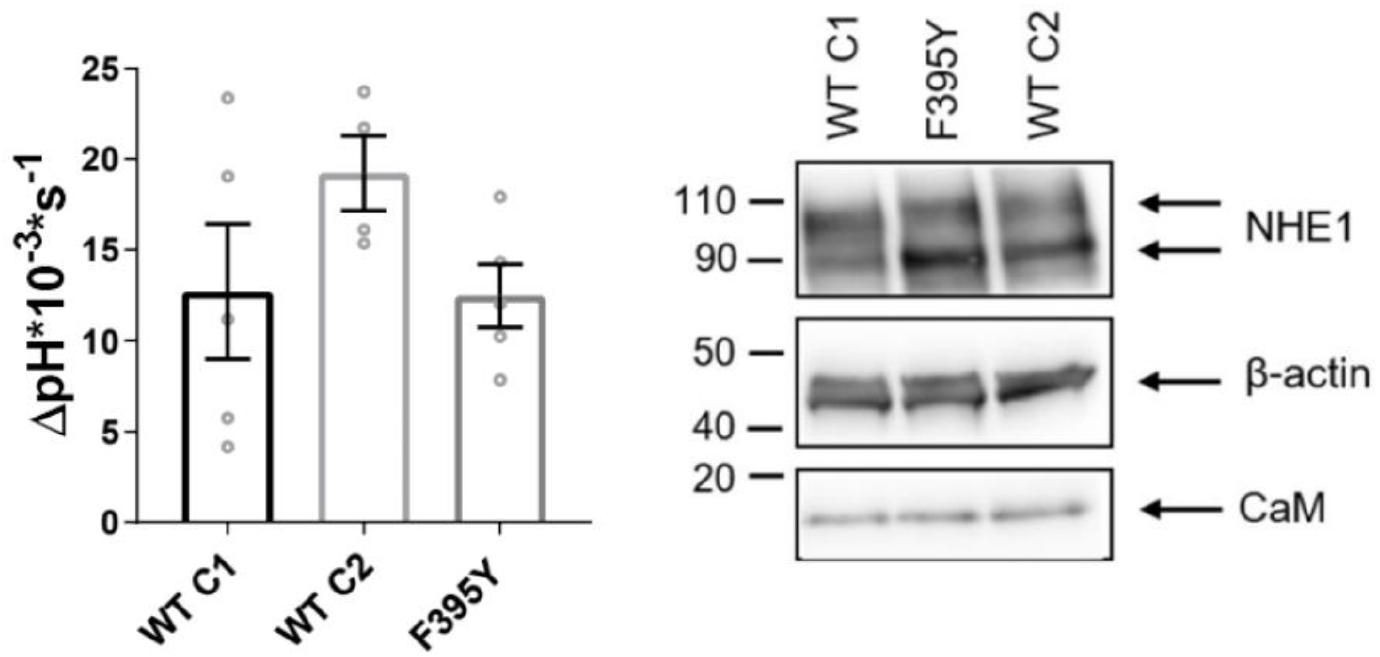
Comparison of pH_*i*_ recovery rate and NHE1 expression levels of three cell lines expressing WT hNHE1 or hNHE1 F395Y. a) Recovery rates as ΔpH*10^−3^*s^−1^ of three PS120 cell lines stably expressing WT hNHE1 (WT C1 and C2) or hNHE1 with a spontaneous F395Y mutation which is present in the cell lines 1K3R1D3E, S648A and S648D. The WT C1 cell line was used as “WT” in all experiments in the main text. pH_*i*_ recovery rate was measured in Ringer solution after an NH_4_Cl-prepulse-induced acid load *(n*=*4-5)*. One-way ANOVA with Dunnett’s post-test comparing cell lines to WT C1 was carried out. b) Representative Western blot of these PS120 cell lines blotted for NHE1 and endogenous CaM. ̫-actin was used as loading control *(n*=*5)*.

**Figure 6 – Figure supplement 1.**
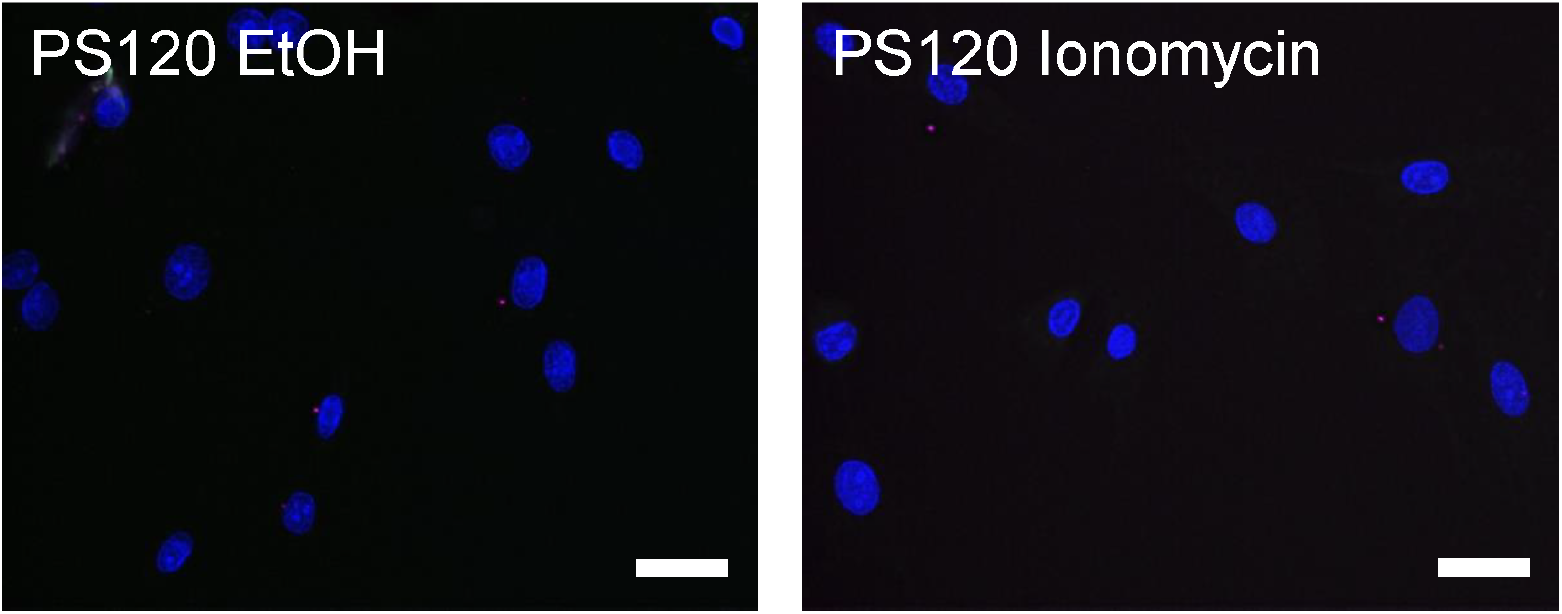
Lack of PLA signal in PS120 cells lacking NHE1. Representative images of PS120 control cells lacking NHE1 and subjected to proximity ligation assay (PLA) with both NHE1 and CaM antibodies, and addition of vehicle (EtOH) or 5 μM ionomycin as indicated *(n*=*4-5)*. Images correspond to those shown in Fig. 6A. The PLA signal is magenta, co-staining for NHE1 is green and nuclei (DAPI) are blue. As seen, negligible PLA signal is observed in cells lacking NHE1, confirming assay specificity. Scale bars represent 20 μm.

**Figure 6 – File supplement 2.**
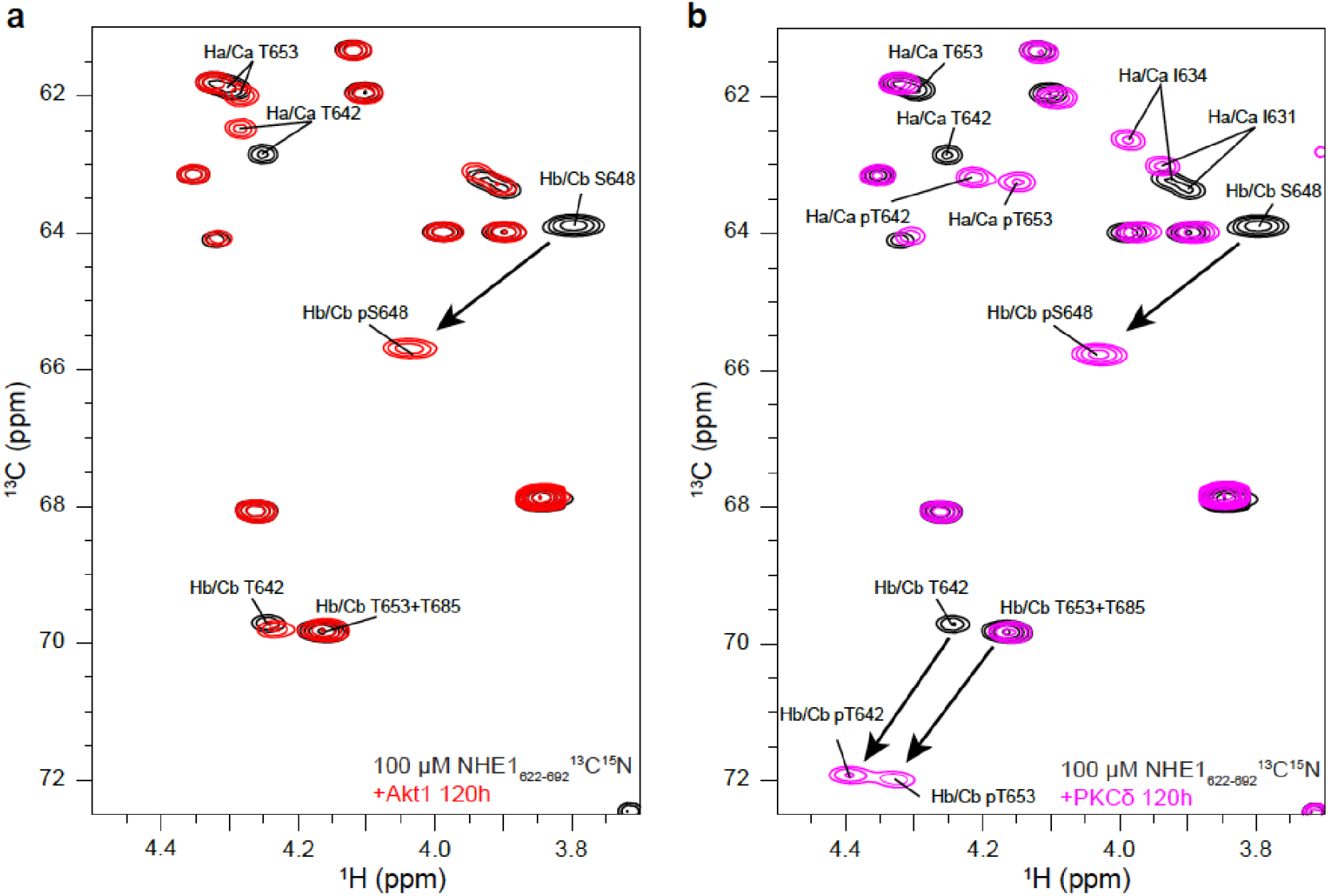
Identification of phosphory/ated residues in NHE1-H1H2. ^13^C, ^1^H HSQC spectra of NHE1-H1H2 before (black) and after 120h of phosphorylation with Akt1 (a, red) or PKCδ (b, pink). The arrows highlight the large shift of the HβCβ peaks upon phosphorylation of residue S648 by (a) Akt1 and residues T642, S648, and T653 by (b) PKCδ.

### Supplementary files not directly related to any main figure

**Supplementary file 1.**
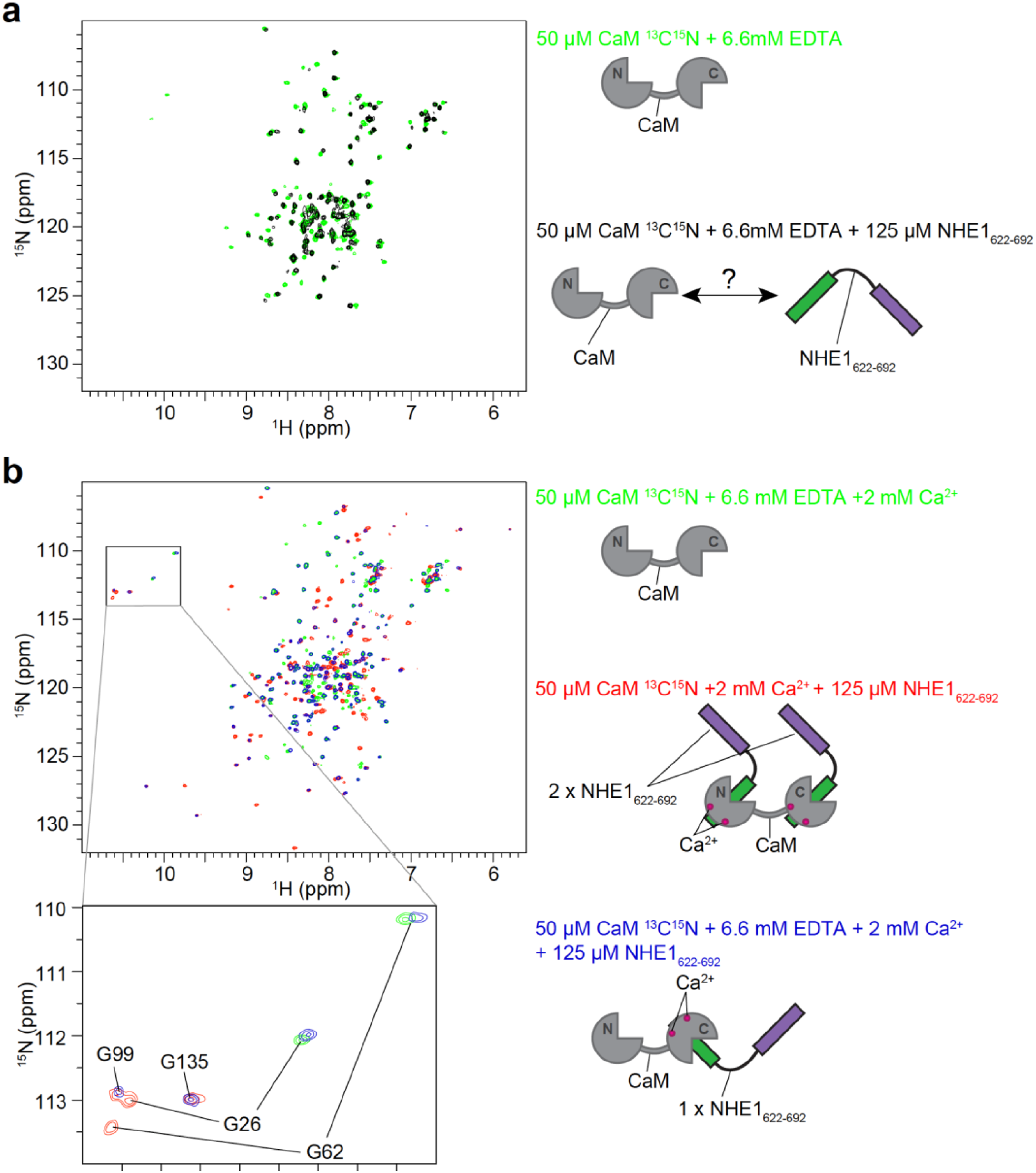
Influence of EDTA on CaM interaction with NHE1_622-692_. (a) In the absence of Ca^2+^, small CSPs and line-broadening of apoCaM were observable after addition of excess NHE1 H1H2 showing that NHE1:CaM complex formation can also occur in the absence of Ca^2+^. (b) In the presence of Ca^2+^ and excess EDTA, free CaM is stripped from Ca^2+^ as can be seen from the similarity of the green ^1^H, ^15^N HSQC spectra in (a) and (b). NHE1 H1H2 increases the Ca^2+^-affinity of the C-lobe of CaM, so that the main observable state in solution has resonances of the N-lobe closely resembling the apo state (G26 and G62), while resonances of the C-lobe resemble the state where the C-lobe is bound to Ca^2+^ and H1 (G99 and G135).

## Data availability

Source data files are provided for Figure 1 (c,d and e) and Figure 6 (h). Resonance assignments of the ternary complex of CaM and two H1 have been deposited in the Biological Magnetic Resonance Bank (BMRB) under ID code 34521. The atomic coordinates for the ternary complex of CaM and two H1 have been deposited in the Protein Data Bank under the ID code 6zbi.

The following datasets were generated:

| Author(s) | Year | Dataset title | Dataset URL | Database and identifier |
| --- | --- | --- | --- | --- |
| Prestel, A, Sjoegaard-Frich, Pedersen, SF, Kragelund, BB | 2020 | $^1\text{H}$, $^{13}\text{C}$ and $^{15}\text{N}$ chemical shift assignments of the | http://www.bmrb.wisc.edu/data_library/summary/i | Biological Magnetic |
|  |  | 2:1 complex between NHE1 and calmodulin | ndex.php?bmrbId=34521 | Resonance Data Bank, 34521 |
| Prestel, A, Sjoegaard-Frich, Pedersen, SF, Kragelund, BB | 2020 | Ternary complex of Calmodulin bound to 2 molecules of NHE1 | http://www.rcsb.org/structure/6zbi | RCSB Protein Data Bank, 6ZBI |

## Code availability

Not applicable.

## References

1. Berchtold, M.W.; Villalobo, A. The many faces of calmodulin in cell proliferation, programmed cell death, autophagy, and cancer. Biochim Biophys Acta 2014, 1843, 398–435, doi:10.1016/j.bbamcr.2013.10.021.

2. Villalobo, A.; Ishida, H.; Vogel, H.J.; Berchtold, M.W. Calmodulin as a protein linker and a regulator of adaptor/scaffold proteins. Biochim Biophys Acta Mol Cell Res 2018, 1865, 507–521, doi:10.1016/j.bbamcr.2017.12.004.

3. Villarroel, A.; Taglialatela, M.; Bernardo-Seisdedos, G.; Alaimo, A.; Agirre, J.; Alberdi, A.; Gomis-Perez, C.; Soldovieri, M.V.; Ambrosino, P.; Malo, C., et al. The ever changing moods of calmodulin: how structural plasticity entails transductional adaptability. J Mol Biol 2014, 426, 2717–2735, doi:10.1016/j.jmb.2014.05.016.

4. Chin, D.; Means, A.R. Calmodulin: a prototypical calcium sensor. Trends Cell Biol 2000, 10, 322–328.

5. Yap, K.L.; Kim, J.; Truong, K.; Sherman, M.; Yuan, T.; Ikura, M. Calmodulin target database. J Struct Funct Genomics 2000, 1, 8–14. DOI: 10.1016/s0962-8924(00)01800-6

6. Tidow, H.; Nissen, P. Structural diversity of calmodulin binding to its target sites. FEBS J 2013, 280, 5551–5565, doi:10.1111/febs.12296.

7. Meador, W.E.; Means, A.R.; Quiocho, F.A. Target enzyme recognition by calmodulin: 2.4 A structure of a calmodulin-peptide complex. Science 1992, 257, 1251–1255. DOI: 10.1126/science.1519061

8. Ikura, M.; Clore, G.M.; Gronenborn, A.M.; Zhu, G.; Klee, C.B.; Bax, A. Solution structure of a calmodulin-target peptide complex by multidimensional NMR. Science 1992, 256, 632–638. DOI: 10.1126/science.1585175

9. Nunomura, W.; Gascard, P.; Wakui, H.; Takakuwa, Y. Phosphatidylinositol-4,5 bisphosphate (PIP(2)) inhibits apo-calmodulin binding to protein 4.1. Biochem Biophys Res Commun 2014, 446, 434–440, doi:10.1016/j.bbrc.2014.02.121.

10. Nunomura, W.; Sasakura, D.; Shiba, K.; Nakamura, S.; Kidokoro, S.; Takakuwa, Y. Structural stabilization of protein 4.1R FERM domain upon binding to apo-calmodulin: novel insights into the biological significance of the calcium-independent binding of calmodulin to protein 4.1R. Biochem J 2011, 440, 367–374, doi:10.1042/BJ20110676.

11. Lee, K.; Kumar, E.A.; Dalby, K.N.; Ghose, R. The role of calcium in the interaction between calmodulin and a minimal functional construct of eukaryotic elongation factor 2 kinase. Protein Sci 2019, 28, 2089–2098, doi:10.1002/pro.3753.

12. Li, L.; Li, Z.; Sacks, D.B. The transcriptional activity of estrogen receptor-alpha is dependent on Ca2+/calmodulin. J Biol Chem 2005, 280, 13097–13104, doi:10.1074/jbc.M410642200.

13. Zhang, Y.; Li, Z.; Sacks, D.B.; Ames, J.B. Structural basis for Ca2+-induced activation and dimerization of estrogen receptor alpha by calmodulin. J Biol Chem 2012, 287, 9336–9344, doi:10.1074/jbc.M111.334797.

14. Whicher, J.R.; MacKinnon, R. Structure of the voltage-gated K(+) channel Eag1 reveals an alternative voltage sensing mechanism. Science 2016, 353, 664–669, doi:10.1126/science.aaf8070.

15. Schonherr, R.; Lober, K.; Heinemann, S.H. Inhibition of human ether a go-go potassium channels by Ca(2+)/calmodulin. EMBO J 2000, 19, 3263–3271, doi:10.1093/emboj/19.13.3263.

16. Girsch, S.J.; Peracchia, C. Calmodulin interacts with a C-terminus peptide from the lens membrane protein MIP26. Curr Eye Res 1991, 10, 839–849. doi: 10.3109/02713689109013880.

17. Orlowski, J.; Grinstein, S. Na+/H+ exchangers. Compr.Physiol 2011, 1, 2083–2100. DOI: 10.1002/cphy.c110020

18. Pedersen, S.F.; Counillon, L. The SLC9A-C Mammalian Na(+)/H(+) Exchanger Family: Molecules, Mechanisms, and Physiology. Physiol Rev 2019, 99, 2015–2113, doi:10.1152/physrev.00028.2018.

19. Stock, C.; Pedersen, S.F. Roles of pH and the Na+/H+ exchanger NHE1 in cancer: From cell biology and animal models to an emerging translational perspective? Semin.Cancer Biol. 2017, 43, 5–16. DOI: 10.1016/j.semcancer.2016.12.001.

20. Cardone, R.A.; Casavola, V.; Reshkin, S.J. The role of disturbed pH dynamics and the Na+/H+ exchanger in metastasis. Nat.Rev.Cancer 2005, 5, 786–795. doi: 10.1038/nrc1713.

21. Imahashi, K.; Mraiche, F.; Steenbergen, C.; Murphy, E.; Fliegel, L. Overexpression of the Na+/H+ exchanger and ischemia-reperfusion injury in the myocardium. Am J Physiol Heart Circ.Physiol 2007, 292, H2237–H2247. DOI: 10.1152/ajpheart.00855.2006.

22. Karmazyn, M.; Gan, X.T.; Humphreys, R.A.; Yoshida, H.; Kusumoto, K. The myocardial Na(+)-H(+) exchange: structure, regulation, and its role in heart disease. Circ.Res. 1999, 85, 777–786. DOI: 10.1161/01.res.85.9.777.

23. Nakamura, T.Y.; Iwata, Y.; Arai, Y.; Komamura, K.; Wakabayashi, S. Activation of Na+/H+ exchanger 1 is sufficient to generate Ca2+ signals that induce cardiac hypertrophy and heart failure. Circ.Res. 2008, 103, 891–899. DOI: 10.1161/CIRCRESAHA.108.175141.

24. Prasad, V.; Chirra, S.; Kohli, R.; Shull, G.E. NHE1 deficiency in liver: implications for non-alcoholic fatty liver disease. Biochem.Biophys.Res.Commun. 2014, 450, 1027–1031.DOI: 10.1016/j.bbrc.2014.06.095

25. Norholm, A.B.; Hendus-Altenburger, R.; Bjerre, G.; Kjaergaard, M.; Pedersen, S.F.; Kragelund, B.B. The intracellular distal tail of the Na+/H+ exchanger NHE1 is intrinsically disordered: implications for NHE1 trafficking. Biochemistry 2011, 50, 3469–3480. DOI: 10.1021/bi1019989.

26. Hendus-Altenburger, R.; Kragelund, B.B.; Pedersen, S.F. Structural Dynamics and Regulation of the Mammalian SLC9A Family of Na(+)/H(+) Exchangers. Curr.Top.Membr. 2014, 73, 69–148. DOI: 10.1016/B978-0-12-800223-0.00002-5

27. Koster, S.; Pavkov-Keller, T.; Kuhlbrandt, W.; Yildiz, O. Structure of human Na+/H+ exchanger NHE1 regulatory region in complex with calmodulin and Ca2+. Journal of Biological Chemistry 2011, 286, 40954–40961. DOI: 10.1074/jbc.M111.286906

28. Bertrand, B.; Wakabayashi, S.; Ikeda, T.; Pouyssegur, J.; Shigekawa, M. The Na+/H+ exchanger isoform 1 (NHE1) is a novel member of the calmodulin-binding proteins. Identification and characterization of calmodulin-binding sites. Journal of Biological Chemistry 1994, 269, 13703–13709. PMID: 8175806

29. Wakabayashi, S.; Bertrand, B.; Ikeda, T.; Pouyssegur, J.; Shigekawa, M. Mutation of calmodulin-binding site renders the Na+/H+ exchanger (NHE1) highly H(+)-sensitive and Ca2+ regulation-defective. Journal of Biological Chemistry 1994, 269, 13710–13715. PMID: 8175807

30. Ikeda, T.; Schmitt, B.; Pouyssegur, J.; Wakabayashi, S.; Shigekawa, M. Identification of cytoplasmic subdomains that control pH-sensing of the Na+/H+ exchanger (NHE1): pH-maintenance, ATP-sensitive, and flexible loop domains. J.Biochem. 1997, 121, 295–303.DOI: 10.1093/oxfordjournals.jbchem.a021586

31. Wakabayashi, S.; Ikeda, T.; Iwamoto, T.; Pouyssegur, J.; Shigekawa, M. Calmodulin-binding autoinhibitory domain controls “pH-sensing” in the Na+/H+ exchanger NHE1 through sequence-specific interaction. Biochemistry 1997, 36, 12854–12861. DOI: 10.1021/bi9715472

32. Li, X.; Prins, D.; Michalak, M.; Fliegel, L. Calmodulin-dependent binding to the NHE1 cytosolic tail mediates activation of the Na+/H+ exchanger by Ca2+ and endothelin. Am J Physiol Cell Physiol 2013, 305, C1161–1169, doi:10.1152/ajpcell.00208.2013.

33. Coaxum, S.D.; Garnovskaya, M.N.; Gooz, M.; Baldys, A.; Raymond, J.R. Epidermal growth factor activates Na(+/)H(+) exchanger in podocytes through a mechanism that involves Janus kinase and calmodulin. Biochim Biophys Acta 2009, 1793, 1174–1181, doi:10.1016/j.bbamcr.2009.03.006.

34. Turner, J.H.; Garnovskaya, M.N.; Coaxum, S.D.; Vlasova, T.M.; Yakutovich, M.; Lefler, D.M.; Raymond, J.R. Ca2+-calmodulin and janus kinase 2 are required for activation of sodium-proton exchange by the Gi-coupled 5-hydroxytryptamine 1a receptor. J Pharmacol.Exp.Ther. 2007, 320, 314–322. DOI: 10.1124/jpet.106.112581

35. Snabaitis, A.K.; Cuello, F.; Avkiran, M. Protein kinase B/Akt phosphorylates and inhibits the cardiac Na+/H+ exchanger NHE1. Circ.Res. 2008, 103, 881–890. DOI: 10.1161/CIRCRESAHA.108.175877

36. Meima, M.E.; Webb, B.A.; Witkowska, H.E.; Barber, D. The Na-H Exchanger NHE1 is an Akt Substrate Necessary for Actin Filament Reorganization by Growth Factors. J Biol Chem. 2009. DOI: 10.1074/jbc.M109.019448

37. Prestel, A.; Bugge, K.; Staby, L.; Hendus-Altenburger, R.; Kragelund, B.B. Characterization of Dynamic IDP Complexes by NMR Spectroscopy. Methods Enzymol 2018, 611, 193–226, doi:10.1016/bs.mie.2018.08.026.

38. Fafournoux, P.; Noel, J.; Pouyssegur, J. Evidence that Na+/H+ exchanger isoforms NHE1 and NHE3 exist as stable dimers in membranes with a high degree of specificity for homodimers. J Biol Chem. 1994, 269, 2589–2596. PMID: 8300588

39. Hisamitsu, T.; Pang, T.; Shigekawa, M.; Wakabayashi, S. Dimeric interaction between the cytoplasmic domains of the Na+/H+ exchanger NHE1 revealed by symmetrical intermolecular cross-linking and selective co-immunoprecipitation. Biochemistry 2004, 43, 11135–11143. DOI: 10.1021/bi049367x

40. Lacroix, J.; Poet, M.; Maehrel, C.; Counillon, L. A mechanism for the activation of the Na/H exchanger NHE-1 by cytoplasmic acidification and mitogens. EMBO Rep. 2004, 5, 91–96. DOI: 10.1038/sj.embor.7400035

41. Hisamitsu, T.; Ammar, Y.B.; Nakamura, T.Y.; Wakabayashi, S. Dimerization Is Crucial for the Function of the Na(+)/H(+) Exchanger NHE1. Biochemistry 2006, 45, 13346–13355. DOI: 10.1021/bi0608616

42. Otsu, K.; Kinsella, J.; Sacktor, B.; Froehlich, J.P. Transient state kinetic evidence for an oligomer in the mechanism of Na+-H+ exchange. Proc.Natl.Acad.Sci.U.S.A 1989, 86, 4818–4822. DOI: 10.1073/pnas.86.13.4818

43. Fuster, D.; Moe, O.W.; Hilgemann, D.W. Steady-state function of the ubiquitous mammalian Na/H exchanger (NHE1) in relation to dimer coupling models with 2Na/2H stoichiometry. J.Gen.Physiol 2008, 132, 465–480. DOI: 10.1085/jgp.200810016

44. Pouyssegur, J.; Sardet, C.; Franchi, A.; L’Allemain, G.; Paris, S. A specific mutation abolishing Na+/H+ antiport activity in hamster fibroblasts precludes growth at neutral and acidic pH. Proc.Natl.Acad.Sci.U.S.A 1984, 81, 4833–4837. DOI: 10.1073/pnas.81.15.4833

45. Amith, S.R.; Wilkinson, J.M.; Fliegel, L. Na+/H+ exchanger NHE1 regulation modulates metastatic potential and epithelial-mesenchymal transition of triple-negative breast cancer cells. Oncotarget. 2016, 7, 21091–21113. DOI: 10.18632/oncotarget.8520

46. Denker, S.P.; Barber, D.L. Cell migration requires both ion translocation and cytoskeletal anchoring by the Na-H exchanger NHE1. J.Cell Biol. 2002, 159, 1087–1096. DOI: 10.1083/jcb.200208050

47. Hendus-Altenburger, R.; Pedraz-Cuesta, E.; Olesen, C.W.; Papaleo, E.; Schnell, J.A.; Hopper, J.T.; Robinson, C.V.; Pedersen, S.F.; Kragelund, B.B. The human Na(+)/H(+) exchanger 1 is a membrane scaffold protein for extracellular signal-regulated kinase 2. BMC.Biol. 2016, 14, 31. DOI: 10.1186/s12915-016-0252-7

48. Hendus-Altenburger, R.; Wang, X.; Sjogaard-Frich, L.M.; Pedraz-Cuesta, E.; Sheftic, S.R.; Bendsoe, A.H.; Page, R.; Kragelund, B.B.; Pedersen, S.F.; Peti, W. Molecular basis for the binding and selective dephosphorylation of Na(+)/H(+) exchanger 1 by calcineurin. Nat Commun 2019, 10, 3489, doi:10.1038/s41467-019-11391-7.

49. Soderberg, O.; Gullberg, M.; Jarvius, M.; Ridderstrale, K.; Leuchowius, K.J.; Jarvius, J.; Wester, K.; Hydbring, P.; Bahram, F.; Larsson, L.G., et al. Direct observation of individual endogenous protein complexes in situ by proximity ligation. Nat.Methods 2006, 3, 995–1000. DOI: 10.1038/nmeth947

50. Edin, S.; Oruganti, S.R.; Grundstrom, C.; Grundstrom, T. Interaction of calmodulin with Bcl10 modulates NF-kappaB activation. Mol Immunol 2010, 47, 2057–2064, doi:10.1016/j.molimm.2010.04.005.

51. Ulke-Lemee, A.; Turner, S.R.; MacDonald, J.A. In situ analysis of smoothelin-like 1 and calmodulin interactions in smooth muscle cells by proximity ligation. J Cell Biochem 2015, 116, 2667–2675, doi:10.1002/jcb.25215.

52. Saucerman, J.J.; Bers, D.M. Calmodulin binding proteins provide domains of local Ca2+ signaling in cardiac myocytes. J Mol Cell Cardiol 2012, 52, 312–316, doi:10.1016/j.yjmcc.2011.06.005.

53. Persechini, A.; Stemmer, P.M. Calmodulin is a limiting factor in the cell. Trends Cardiovasc Med 2002, 12, 32–37, doi:10.1016/s1050-1738(01)00144-x.

54. Mori, M.X.; Erickson, M.G.; Yue, D.T. Functional stoichiometry and local enrichment of calmodulin interacting with Ca2+ channels. Science 2004, 304, 432–435, doi:10.1126/science.1093490.

55. Li, Z.; Zhang, Y.; Hedman, A.C.; Ames, J.B.; Sacks, D.B. Calmodulin Lobes Facilitate Dimerization and Activation of Estrogen Receptor-alpha. J Biol Chem 2017, 292, 4614–4622, doi:10.1074/jbc.M116.754804.

56. Schumacher, M.A.; Rivard, A.F.; Bachinger, H.P.; Adelman, J.P. Structure of the gating domain of a Ca2+-activated K+ channel complexed with Ca2+/calmodulin. Nature 2001, 410, 1120–1124, doi:10.1038/35074145.

57. Lee, C.H.; MacKinnon, R. Activation mechanism of a human SK-calmodulin channel complex elucidated by cryo-EM structures. Science 2018, 360, 508–513, doi:10.1126/science.aas9466.

58. Barros, F.; Pardo, L.A.; Dominguez, P.; Sierra, L.M.; de la Pena, P. New Structures and Gating of Voltage-Dependent Potassium (Kv) Channels and Their Relatives: A Multi-Domain and Dynamic Question. Int J Mol Sci 2019, 20, doi:10.3390/ijms20020248.

59. Hisamitsu, T.; Nakamura, T.Y.; Wakabayashi, S. Na(+)/H(+) exchanger 1 directly binds to calcineurin A and activates downstream NFAT signaling, leading to cardiomyocyte hypertrophy. Mol.Cell Biol. 2012, 32, 3265–3280. DOI: 10.1128/MCB.00145-12

60. Aharonovitz, O.; Zaun, H.C.; Balla, T.; York, J.D.; Orlowski, J.; Grinstein, S. Intracellular pH regulation by Na(+)/H(+) exchange requires phosphatidylinositol 4,5-bisphosphate. J.Cell Biol. 2000, 150, 213–224. DOI: 10.1083/jcb.150.1.213

61. Shimada-Shimizu, N.; Hisamitsu, T.; Nakamura, T.Y.; Hirayama, N.; Wakabayashi, S. Na+/H+ exchanger 1 is regulated via its lipid-interacting domain, which functions as a molecular switch: a pharmacological approach using indolocarbazole compounds. Mol.Pharmacol. 2014, 85, 18–28. DOI: 10.1124/mol.113.089268

62. Monteiro, M.E.; Sarmento, M.J.; Fernandes, F. Role of calcium in membrane interactions by PI(4,5)P(2)-binding proteins. Biochem Soc Trans 2014, 42, 1441–1446, doi:10.1042/BST20140149.

63. Cao, C.; Zakharian, E.; Borbiro, I.; Rohacs, T. Interplay between calmodulin and phosphatidylinositol 4,5-bisphosphate in Ca2+-induced inactivation of transient receptor potential vanilloid 6 channels. J Biol Chem 2013, 288, 5278–5290, doi:10.1074/jbc.M112.409482.

64. Hendus-Altenburger, R.V., J.; Pedersen, E.; Luchini, A.; Araya-Secchi, R.; Bendsoe, A.; Prestel, A.; Cardenas, M.; Pedraz-Cuesta E.; Arleth, L.; Pedersen S.F.; Kragelund B.B. The lipid-binding domain of human Na+/H+ exchanger 1 forms a 4 helical lipid-protein co-structure essential for activity: a new 5 principle for membrane protein regulation via disordered domains. in review 2020.

65. Rust, H.L.; Thompson, P.R. Kinase consensus sequences: a breeding ground for crosstalk. ACS Chem Biol 2011, 6, 881–892, doi:10.1021/cb200171d.

66. Wallert, M.A.; Hammes, D.; Nguyen, T.; Kiefer, L.; Berthelsen, N.; Kern, A.; Anderson-Tiege, K.; Shabb, J.B.; Muhonen, W.W.; Grove, B.D., et al. RhoA Kinase (Rock) and p90 Ribosomal S6 Kinase (p90Rsk) phosphorylation of the sodium hydrogen exchanger (NHE1) is required for lysophosphatidic acid-induced transport, cytoskeletal organization and migration. Cell Signal. 2015, 27, 498–509. DOI: 10.1016/j.cellsig.2015.01.002

67. Li, X.; Augustine, A.; Sun, D.; Li, L.; Fliegel, L. Activation of the Na+/H+ exchanger in isolated cardiomyocytes through beta-Raf dependent pathways. Role of Thr653 of the cytosolic tail. J Mol.Cell Cardiol. 2016, 99, 65–75. DOI: 10.1016/j.yjmcc.2016.08.014

68. Boyarsky, G.; Ganz, M.B.; Sterzel, R.B.; Boron, W.F. pH regulation in single glomerular mesangial cells. II. Na+-dependent and -independent Cl(−)-HCO3(−) exchangers. Am.J Physiol 1988, 255, C857–C869. DOI: 10.1152/ajpcell.1988.255.6.C857

69. Pedersen, S.F.; Jorgensen, N.K.; Damgaard, I.; Schousboe, A.; Hoffmann, E.K. Mechanisms of pHi regulation studied in individual neurons cultured from mouse cerebral cortex. J.Neurosci.Res. 1998, 51, 431–441. DOI: 10.1002/(SICI)1097-4547(19980215)51:4<431::AID-JNR3>3.0.CO;2-D

70. Delaglio, F.; Grzesiek, S.; Vuister, G.W.; Zhu, G.; Pfeifer, J.; Bax, A. NMRPipe: a multidimensional spectral processing system based on UNIX pipes. J.Biomol.NMR. 1995, 6, 277–293. DOI: 10.1007/BF00197809

71. Kazimierczuk, K.; Orekhov, V.Y. Accelerated NMR spectroscopy by using compressed sensing. Angew Chem Int Ed Engl 2011, 50, 5556–5559, doi:10.1002/anie.201100370.

72. Vranken, W.F.; Boucher, W.; Stevens, T.J.; Fogh, R.H.; Pajon, A.; Llinas, M.; Ulrich, E.L.; Markley, J.L.; Ionides, J.; Laue, E.D. The CCPN data model for NMR spectroscopy: development of a software pipeline. Proteins. 2005, 59, 687–696. DOI: 10.1002/prot.20449

73. Waudby, C.A.; Ramos, A.; Cabrita, L.D.; Christodoulou, J. Two-Dimensional NMR Lineshape Analysis. Scientific Reports 2016, 6, 24826, doi:10.1038/srep24826.

74. Shen, Y.; Bax, A. Protein backbone and sidechain torsion angles predicted from NMR chemical shifts using artificial neural networks. Journal of biomolecular NMR 2013, 56, 227–241, doi:10.1007/s10858-013-9741-y.

75. Güntert, P.; Buchner, L. Combined automated NOE assignment and structure calculation with CYANA. Journal of biomolecular NMR 2015, 62, 453–471, doi:10.1007/s10858-015-9924-9.

76. Chattopadhyaya, R.; Meador, W.E.; Means, A.R.; Quiocho, F.A. Calmodulin structure refined at 1.7 A resolution. J Mol Biol 1992, 228, 1177–1192, doi:10.1016/0022-2836(92)90324-d.

77. Tian, Y.; Schwieters, C.D.; Opella, S.J.; Marassi, F.M. A practical implicit solvent potential for NMR structure calculation. Journal of magnetic resonance (San Diego, Calif. : 1997) 2014, 243, 54–64, doi:10.1016/j.jmr.2014.03.011.

78. Bermejo, G.A.; Schwieters, C.D. Protein Structure Elucidation from NMR Data with the Program Xplor-NIH. Methods Mol Biol 2018, 1688, 311–340, doi:10.1007/978-1-4939-7386-6_14.

79. Munoz, V.; Serrano, L. Development of the multiple sequence approximation within the AGADIR model of alpha-helix formation: comparison with Zimm-Bragg and Lifson-Roig formalisms. Biopolymers 1997, 41, 495–509, doi:10.1002/(SICI)1097-0282(19970415)41:5<495::AID-BIP2>3.0.CO;2-H.

